# LITESEC-T3SS - Light-controlled protein delivery into eukaryotic cells with high spatial and temporal resolution

**DOI:** 10.1101/807461

**Authors:** Florian Lindner, Bailey Milne-Davies, Katja Langenfeld, Andreas Diepold

## Abstract

Many bacteria employ a type III secretion system (T3SS), also called injectisome, to translocate proteins into eukaryotic host cells through a hollow extracellular needle. The system can efficiently transport heterologous cargo, which makes it a uniquely suited tool for the translocation of proteins into eukaryotic cells. However, the injectisome indiscriminately injects proteins into any adjoining eukaryotic cell, and this lack of target specificity currently limits its application in biotechnology and healthcare. In this study, we exploit the dynamic nature of the T3SS to control protein secretion and translocation into eukaryotic cells by light. By combining optogenetic interaction switches with the dynamic cytosolic T3SS component SctQ, the cytosolic availability of SctQ and in consequence T3SS-dependent effector secretion can be regulated by external light. The resulting system, which we call LITESEC-T3SS (**L**ight-**i**nduced **t**ranslocation of **e**ffectors through **s**equestration of **e**ndogenous **c**omponents of the **T3SS**), allows rapid, specific, and reversible activation or deactivation of the T3SS upon illumination. We demonstrate the application of the system for light-regulated translocation of a heterologous reporter protein into cultured eukaryotic cells. LITESEC-T3SS represents a new method to achieve unparalleled spatial and temporal resolution for the controlled protein translocation into eukaryotic host cells.

## Introduction

### The bacterial type III secretion injectisome

The injectisome is a bacterial nanomachine capable of translocating proteins into eukaryotic host cells in a one-step export mechanism^1^,^2^. The core components of the injectisome, or type III secretion system (T3SS)^§^ are shared with the bacterial flagellum^3^,^4^. The injectisome consists of (i) an extracellular needle formed by helical polymerization of a small protein and terminated by a pentameric tip structure, (ii) a series of membrane rings that span both bacterial membranes and embed (iii) the export apparatus, formed by five highly conserved hydrophobic proteins thought to gate the export process, and (iv) a set of essential cytosolic components, which cooperate in substrate selection and export (Fig. 1A).

**Fig. 1:**
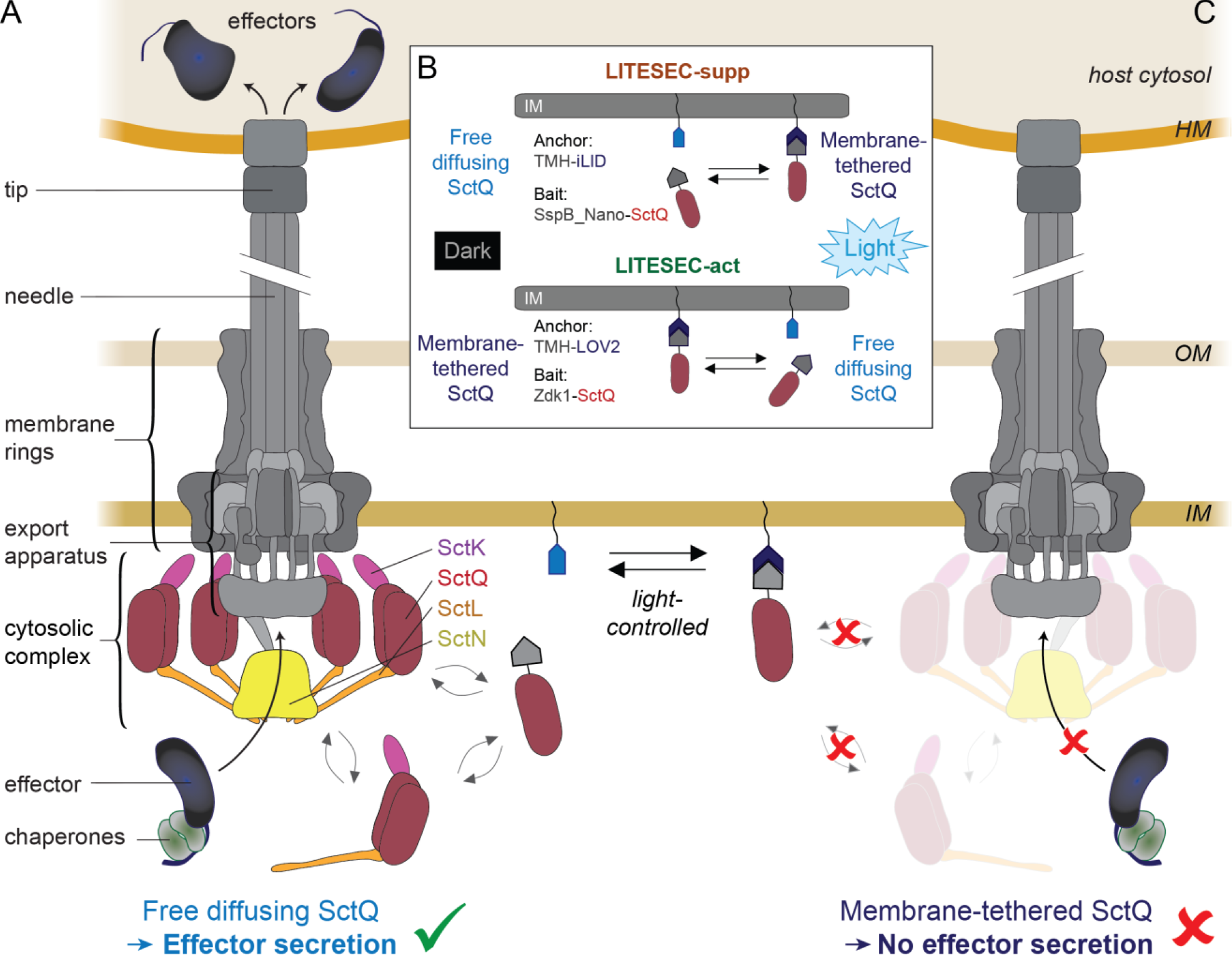
Working principle of the LITESEC systems – light-controlled activation and deactivation of protein translocation by the type III secretion system. (**A**) Schematic representation of the active T3SS injectisome (modified from ref. 39). Left side, main substructures; right side, dynamic cytosolic T3SS components. Effector translocation by the T3SS is licensed by the functional interaction of the unbound bait-SctQ fusion with the T3SS. (**B**) Different states of the bait and anchor proteins in dark and light conditions. In the LITESEC-supp system (top), the bait protein, a fusion of the smaller interaction switch domain SspB_Nano and the essential T3SS component SctQ, is tethered to the inner membrane (IM) by a membrane anchor, a fusion of a transmembrane helix (TMH) and the larger interaction switch domain, iLID, in the light, and gets released in the dark. Conversely, in the LITESEC-act system (bottom), the bait protein, a fusion of the smaller interaction switch domain, Zdk1, and the essential T3SS component SctQ, is tethered to the membrane anchor, a TMH fusion of the larger interaction switch domain, LOV2, in the dark, and gets released by illumination. (**C**) Sequestration of the bait-SctQ fusion protein to the membrane prevents effector secretion. HM, host membrane; OM, bacterial outer membrane; IM, bacterial inner membrane.

The injectisome is an essential virulence factor for many pathogenic Gram-negative bacteria, including *Salmonella*, *Shigella*, pathogenic *Escherichia coli*, and *Yersinia*^5^. It is usually assembled upon entry into a host organism, but remains inactive until contact to a host cell has been established. At this point, the injectisome exports two translocator proteins that form a pore in the host membrane, and a pool of so-called T3SS effector proteins that are translocated into the host cell.

The Gram-negative enterobacterium *Y. enterocolitica* uses the T3SS to translocate six Yop (*Yersinia* outer protein) effector proteins into phagocytes, which prevent phagocytosis and block pro-inflammatory signaling^7^. In this study, we use the *Y. enterocolitica* strain IML421asd (∆HOPEMTasd)^8^, where these six virulence effectors have been deleted, and which is additionally auxotrophic for the cell wall component diaminopimelic acid. The strain is therefore non-pathogenic, but possesses a functional T3SS. Secretion of effector proteins can be triggered *in vivo* by host cell contact or *in vitro* by low Ca^2+^ levels in the medium^9^.

### The T3SS as a protein translocation device

Being a machinery that evolved to efficiently translocate proteins into eukaryotic cells, the T3SS has been successfully used to deliver protein cargo into a wide variety of eukaryotic target cells for different purposes such as vaccination, immunotherapy, and gene editing (reviewed in ref. ^10^). Export through the T3SS is fast and efficient: More than 10^6^ effectors can be translocated into a single host cell at rates of several hundred effectors per second for one injectisome^11–14^. Short N-terminal secretion signals mark cargo proteins for delivery by the T3SS^15^,^16^. The size and structure of the cargo proteins can influence translocation rates, and very large or stably folded proteins (such as GFP or dihydrofolate reductase) are exported at a lower rate. However, most cargoes, including large proteins with molecular weights above 60 kDa, can be exported by the T3SS^14,17,18^. Protein translocation into host cells can be titrated by adjusting the expression level and multiplicity of infection (ratio of bacteria and host cells). Within the host, the T3SS secretion signal can be removed by site-specific proteases or cleavage at the C-terminus of a ubiquitin domain by the native host cell machinery, and subcellular localization can be influenced using nanobodies co-translocated by the T3SS^14,19^. Taken together, these properties make the T3SS an efficient and versatile tool for protein delivery into eukaryotic cells^10,14^.

T3SS inject effector proteins into any eukaryotic host cell as soon as they are in contact. Lack of target specificity is therefore a main obstacle in the further development and application of T3SS-based protein delivery systems^20,21^.

### Dynamics of the cytosolic components of the T3SS and its link to effector secretion

Four soluble cytosolic components of the T3SS (SctK, L, Q, N) form an interdependent complex at the proximal interface of the injectisome^22–29^ (Fig. 1A). As these proteins interact with effectors and their chaperones with a graded affinity matching the export order of the effectors, they were termed “sorting platform”^30^. Our group recently discovered that the sorting platform proteins of the *Y. enterocolitica* T3SS constantly exchange between the injectisome and a cytosolic pool (Fig. 1A), and that this exchange is linked to protein secretion by the T3SS^25,31^. We rationalized that the constant shuttling of these essential T3SS components should allow to control T3SS activity through reversible sequestration of one of the cytosolic proteins, thereby establishing a completely new way of regulating the T3SS.

### Optogenetic control of protein interactions

Optogenetics combines optical and genetic methods to precisely and reversibly control gain or loss of protein function in living cells or tissues. It allows fast (within milliseconds) and specific (to single proteins) control of defined events in biological systems^32^, giving optogenetic approaches an advantage over knockdown, overexpression, or mutant strain analysis, which often display slower activation and a broader effect^33^. Optogenetic protein interaction switches use light-induced conformational changes of specific proteins, often light-oxygen-voltage (LOV) domain proteins, to control protein-protein interactions by light^34,35^. They usually consist of homo-or hetero-dimers whose affinities are strongly altered upon irradiation by light of a certain wavelength. Mutations of specific amino acids in the optogenetic interaction domains can modulate binding affinities and the corresponding dissociation or return rates from a few seconds to several minutes^35,36^.

Optogenetic interaction switches were established and have mainly been studied in eukaryotic cells^37^. In this work, we therefore tested the applicability of two different optogenetic interaction switches in bacteria: (i) The LOVTRAP system (LOV), which consists of the two interacting proteins LOV2 (a photo sensor domain from *Avena sativa* phototropin 1) and Zdk1 (Z subunit of the protein A), that bind to each other in the dark. Upon irradiation with blue light, LOV2 undergoes a conformational change and Zdk1 is released^35^. (ii) The iLID system, which employs the interaction of iLID, derived from a LOV2 domain from *Avena sativa* phototropin 1, with a smaller binding partner, SspB_Nano. The iLID system has a low binding affinity in the dark and a high affinity upon irradiation with blue light^34,36^. LOV and iLID systems therefore react to light in opposite directions, which allows to specifically release a bait protein (and, subsequently, to activate processes that require its presence) in the dark or upon illumination, respectively.

To establish the use of optogenetic interaction switches in bacteria, we first assessed the effect of illumination on the different switches by light microscopy, using fluorescently labeled bait proteins. Next, we applied the switches to control the availability of the essential cytosolic T3SS component SctQ and, in consequence, secretion of cargo proteins through the T3SS, by light. We optimized the systems by defining suitable versions of the switches and adjusting the expression ratio of anchor and bait proteins. As proof of concept, we show the light-dependent translocation of a heterologous cargo protein into eukaryotic host cells. The successful development of the LITESEC system presents a blueprint for the application of optogenetic interaction switches in prokaryotes, and opens widespread opportunities for using the T3SS as a specific and precisely controllable tool to deliver proteins of interest into eukaryotic cells.

## Results

### Controlling protein secretion and translocation by the T3SS with light

To establish a method to control protein translocation by the T3SS, we took advantage of our recent finding that some essential cytosolic T3SS components constantly exchange between the cytosol and the injectisome^25,31^. We combined one of these components, SctQ, with one partner domain of an optogenetic interaction switch, and targeted the other partner domain to the bacterial inner membrane (IM) by adding an N-terminal transmembrane helix. This allowed to control SctQ availability in the cytosol, and therefore T3SS-based protein export and translocation into host cells, by light. To be able to control T3SS activity in both directions, we developed two complementary systems:

A. LITESEC-supp, a system that confers suppression of T3SS-dependent protein translocation by blue light illumination
B. LITESEC-act, a system that confers activation of T3SS-dependent protein translocation by blue light illumination

Both systems rely on two interaction partners which we have engineered:

i. A membrane-bound **anchor protein**, which is a fusion between the N-terminal transmembrane helix (TMH) of a well-characterized transmembrane protein, *Escherichia coli* TatA, extended by two amino acids (Val-Leu) for more stable insertion in the IM, a Flag peptide for detection and spacing, and the larger domain of the respective optogenetic interaction switches, iLID (for LITESEC-supp) or LOV2 (for LITESEC-act). The resulting fusion proteins, **TMH-iLID / TMH-LOV2**, are expressed from a plasmid.
ii. A **bait protein**, which consists of a fusion between the essential cytosolic T3SS component SctQ and the smaller domain of the interaction switches, SspB_Nano (LITESEC-supp) / Zdk1 (LITESEC-act). The resulting fusion proteins, **SspB_Nano-SctQ / Zdk1-SctQ**, replace the wild-type SctQ protein on the *Y. enterocolitica* virulence plasmid by allelic exchange of the genes^38^.

Co-expression of both proteins provides the basis for light-controlled protein translocation by the T3SS (Fig. 1). For the iLID-based LITESEC-supp system, the bait protein is tethered to the membrane anchor in the light, and SctQ is therefore not available to interact with the T3SS (Fig. 1B). As SctQ is essential for the function of the T3SS, protein secretion by the T3SS is prevented (Fig. 1C). In the dark, the bait protein is not bound to the membrane, and can therefore functionally interact with the T3SS, allowing protein secretion by the T3SS (Fig. 1A). Conversely, in the LOV-based LITESEC-act system, the bait protein is released from the membrane upon irradiation with blue light, licensing protein secretion by the T3SS (Fig. 1).

### Characterization of optogenetic sequestration systems in *Y. enterocolitica*

To assess the function and efficiency of the used optogenetic interaction switches as sequestration systems in prokaryotes, and to monitor their dynamics, we visualized the components of iLID- and LOV-based sequestration systems^34,35^ in live *Y. enterocolitica* by time-lapse fluorescence microscopy. We coexpressed the anchor protein with a version of the corresponding bait protein where SctQ was replaced by mCherry to allow for a characterization of the switch by fluorescence microscopy. Initially, we confirmed that mCherry fused to the membrane anchor showed a strict membrane localization (Suppl. Fig. 1), indicating a stable fusion and a functional TMH motif. Next, the localization of mCherry-bait fusions was determined by fluorescence microscopy in live *Y. enterocolitica* expressing the corresponding unlabeled anchor proteins (Suppl. Table 1). Bacteria were grown in the dark and the distribution of the bait proteins was monitored before and after a short pulse of blue light (Fig. 2AB). To quantify the change of the normalized fluorescence signal across the bacterial cells, line scans were performed (Fig. 2CD). For the iLID system, the fluorescence signal of the bait-mCherry was cytosolic in the pre-activated state. After activation of the interaction switch with blue light, the fluorescence signal partly shifted to the membrane (Fig. 2A) and returned to the cytosol within the next minutes (Fig. 2C). In contrast, for the LOV-based sequestration system, the fluorescence signal of the bait-mCherry was mainly membrane localized in the pre-activated state. Activation with blue light led to only a minor relocalization of the signal from the membranes to the cytosol (Fig. 2BD), suggesting that the majority of bait protein remained bound to the anchor even after illumination.

**Fig. 2:**
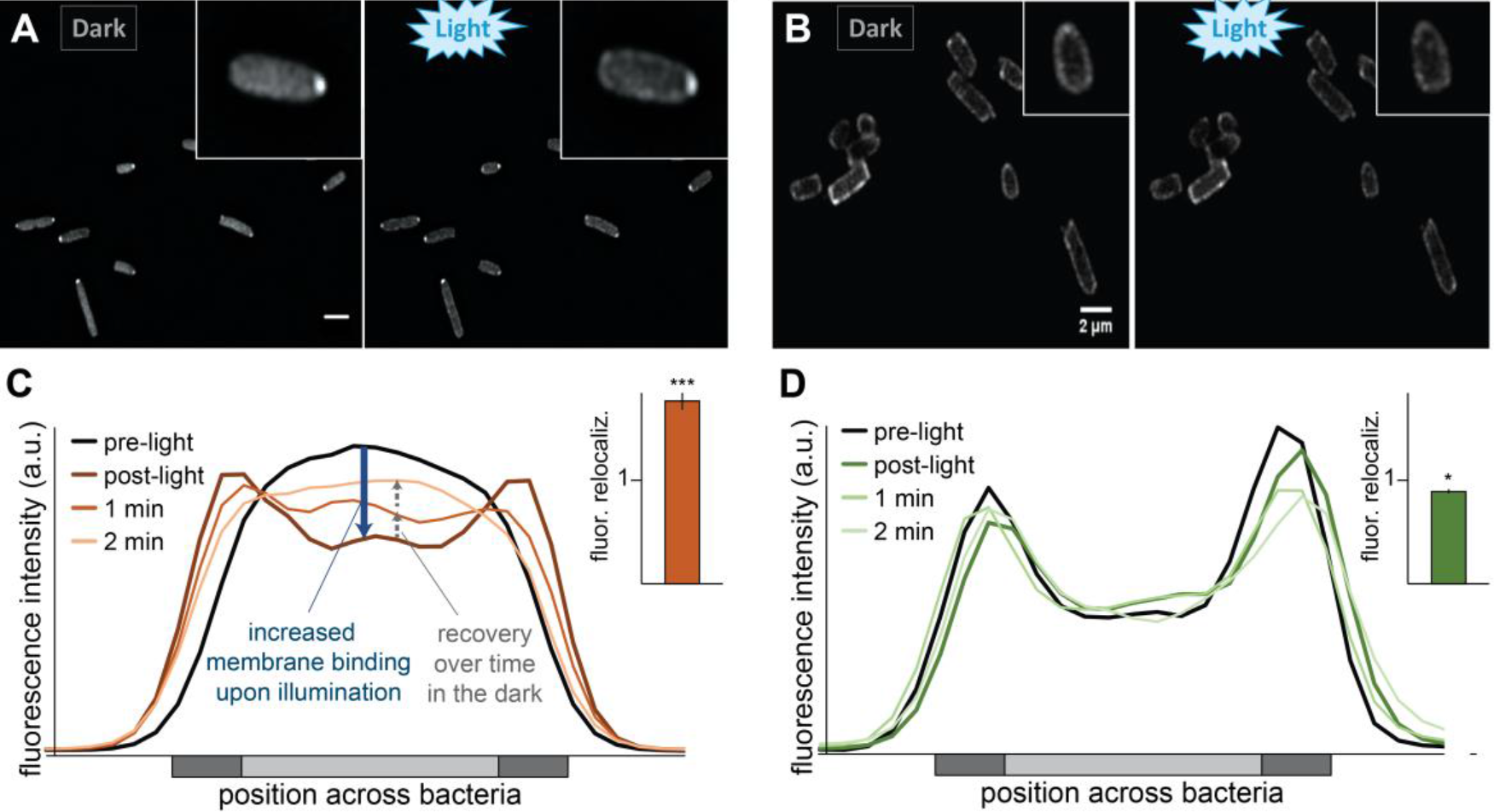
Activation and recovery kinetics of optogenetic sequestration systems.

(**A**/**B**) Fluorescence micrographs of mCherry-labeled bait proteins in the iLID-based (A) and LOV-based (B) sequestration systems, before (left) and directly after (right) illumination with blue light. (**C**/**D**) Representative fluorescence signal quantification across bacteria over time in the iLID-based (C) and LOV-based (D) sequestration systems; dark grey: membrane, light grey: cytosol. Insets: Fluorescence relocalization factor (fluor. reloc. = R_post-light_/R_pre-light_, where R represents the ratio of fluorescence intensities at the membrane and in the cytosol, before and after illumination, respectively), based on 121-131 line scans across five cells per strain and time point. Error bars represent the standard deviation, ^*^, p<0.05; ^***^, p<0.001 against no relocalization in a two-tailed homoscedastic t-test.

### Development and characterization of LITESEC strains

For the development of the LITESEC strains, we replaced SctQ with the bait fusion proteins Zdk1-SctQ or SspB_Nano-SctQ at its native genetic location via allelic exchange. We confirmed the stability of the fusion proteins in the LITESEC strains by Western blot (Suppl. Fig. 2). Protein secretion in wild-type *Y. enterocolitica* was not influenced by the used illumination (Suppl. Fig. 3A), and the blue light had no influence on growth of *Y. enterocolitica* (Suppl. Fig. 3B).

### Inhibition of protein secretion by illumination in the LITESEC-supp system

Can we use LITESEC to control T3SS secretion by light? We first tested the LITESEC-supp1 system, designed to suppress T3SS protein secretion upon illumination (Table 1), in an *in vitro* protein secretion assay under conditions that usually lead to effector secretion (presence of 5 mM EGTA in the medium)^9^. The control strain lacking the membrane anchor secreted effectors irrespective of the illumination (Fig. 3A, lanes 4, 5), confirming the functionality of the used SctQ fusion protein. Strikingly, the LITESEC-supp1 system showed a high level of secretion when grown in the dark, but strongly reduced secretion when grown under blue light (Fig. 3A, lanes 6, 7). To quantify the difference of secretion under light and dark conditions, we define the light/dark secretion ratio (L/D ratio) as the ratio of secretion efficiency under light and dark conditions. For the LITESEC-supp1 system, the L/D ratio was 0.28, with normalized secretion efficiencies of 23.5 ± 8.1% and 85.1 ± 5.1% in light and dark conditions, respectively.

**Table 1.** Schematic overview of the LITESEC systems and their optogenetic components. Overview of the interaction partners and their properties. All bait proteins are expressed from their native genetic locus. TMH, extended transmembrane helix (see material and methods for details).

| Optogenetic T3SS control system | Anchor (*plasmid*) | Bait (*endogenous expression*) | Properties |
| --- | --- | --- | --- |
| LITESEC-supp1<br>LITESEC-supp2 | TMH-FLAG-iLID (*pBAD*)<br>TMH-FLAG-iLID (*pACYC184*) | SspB_Nano-SctQ | Suppression of T3SS-based protein secretion upon illumination by membrane sequestration of essential cytosolic T3SS component |
| LITESEC-act1<br>LITESEC-act2<br>LITESEC-act3 | TMH-FLAG-LOV2 (*pBAD*)<br>TMH-FLAG-LOV2<sub>V416L</sub> (*pBAD*)<br>TMH-FLAG-LOV2<sub>V416L</sub> (*pACYC184*) | Zdk1-SctQ | Activation of T3SS-based protein secretion upon illumination by release of essential cytosolic T3SS component |

**Fig. 3:**
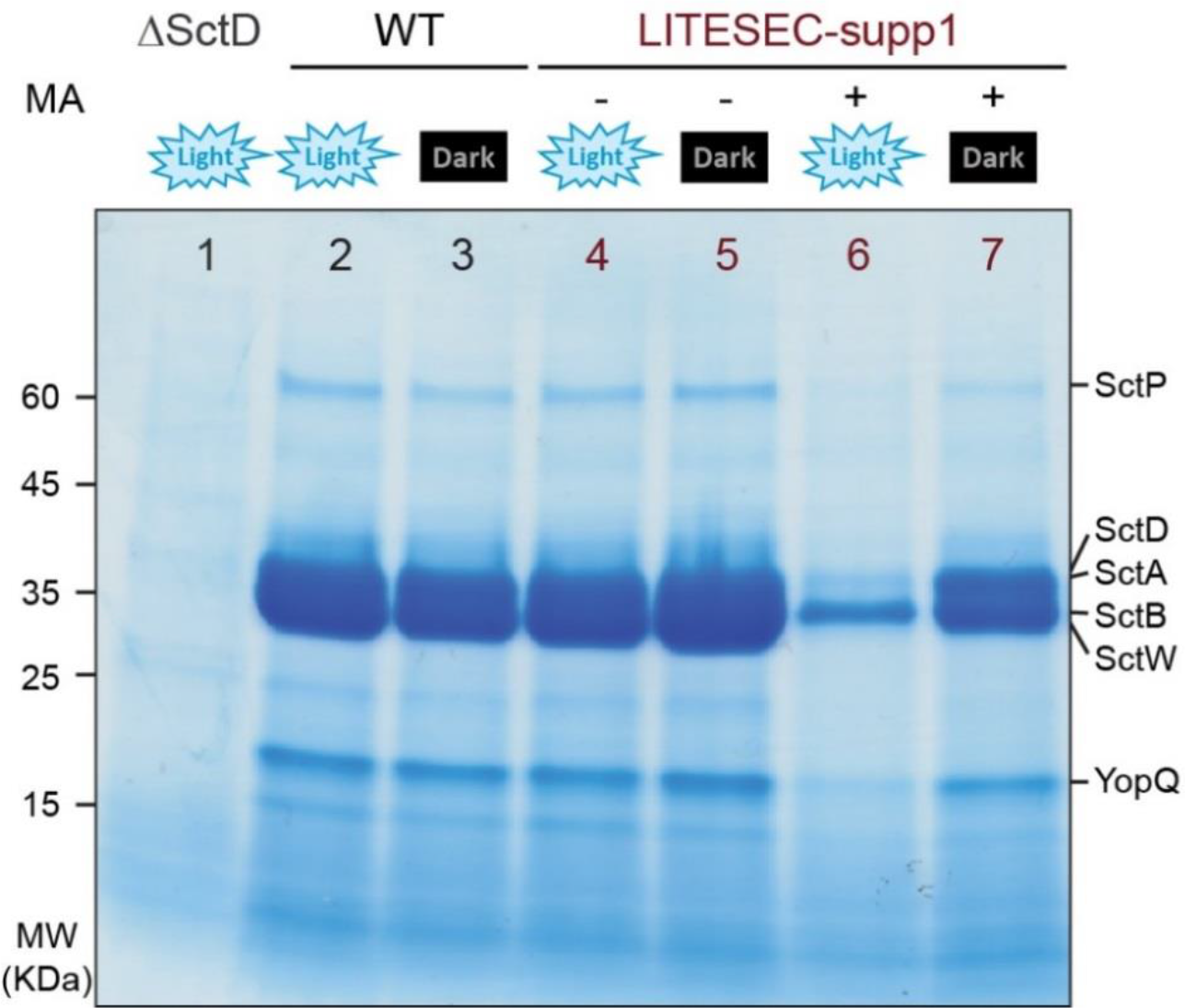
Secretion of effector proteins by the type III secretion system can be controlled by light. *In vitro* secretion assay showing light-dependent export of native T3SS substrates (indicated on the right) in the LITESEC-supp1 strain. Proteins secreted by 3^*^10^9^ bacteria during a 180 min incubation period were precipitated and analyzed by SDS-PAGE. A strain lacking the membrane anchor (MA), the wild-type strain ∆HOPEMTasd and the T3SS-negative strain ∆SctD were used as controls. MW, molecular weight in kDa.

### Improved functionality of the LITESEC-act system with a mutated anchor (V416L)

We next tested the LITESEC-act1 system, designed for induction of secretion by blue light illumination (Table 1), and detected only a very weak activation of protein export under light conditions (Fig. 4, lanes 1-2). Based on the fact that secretion was wild-type-like in the absence of the membrane anchor (Fig. 4, lane 7), and the results of the earlier sequestration experiment (Fig. 2BD), we concluded that bait and anchor interact too strongly in the LITESEC-act1 system. Therefore, we constructed and tested additional versions of the system, using the mutated anchor version V416L, which displays a weaker affinity to the bait^35^. We hypothesized that a lower anchor/bait expression ratio could additionally lead to more efficient release of the bait and activation of T3SS secretion upon illumination, and expressed the V416L version of the anchor both from the medium-high copy pBAD expression vector used previously (LITESEC-act2), and a constitutive low-copy vector, pACYC184 (LITESEC-act3). The response of the resulting LITESEC systems (Table 1) to light was tested in an *in vitro* secretion assay. LITESEC-act2 showed significant induction of protein secretion in the light, compared to dark conditions (L/D ratio 2.16, Fig. 4, lanes 3-4). Even more markedly, LITESEC-act3 allowed an almost complete activation of secretion upon illumination (L/D ratio 4.18, Fig. 4, lanes 5-6). Both new strains retained the low level of export in the dark. We also expressed the anchor for the LITESEC-supp system from pACYC184. The resulting LITESEC-supp2 system showed efficient secretion in the dark and strong suppression of secretion upon illumination (L/D ratio 0.26), comparable with the LITESEC-supp1 system (Fig. 4, lanes 8-11).

**Fig. 4:**
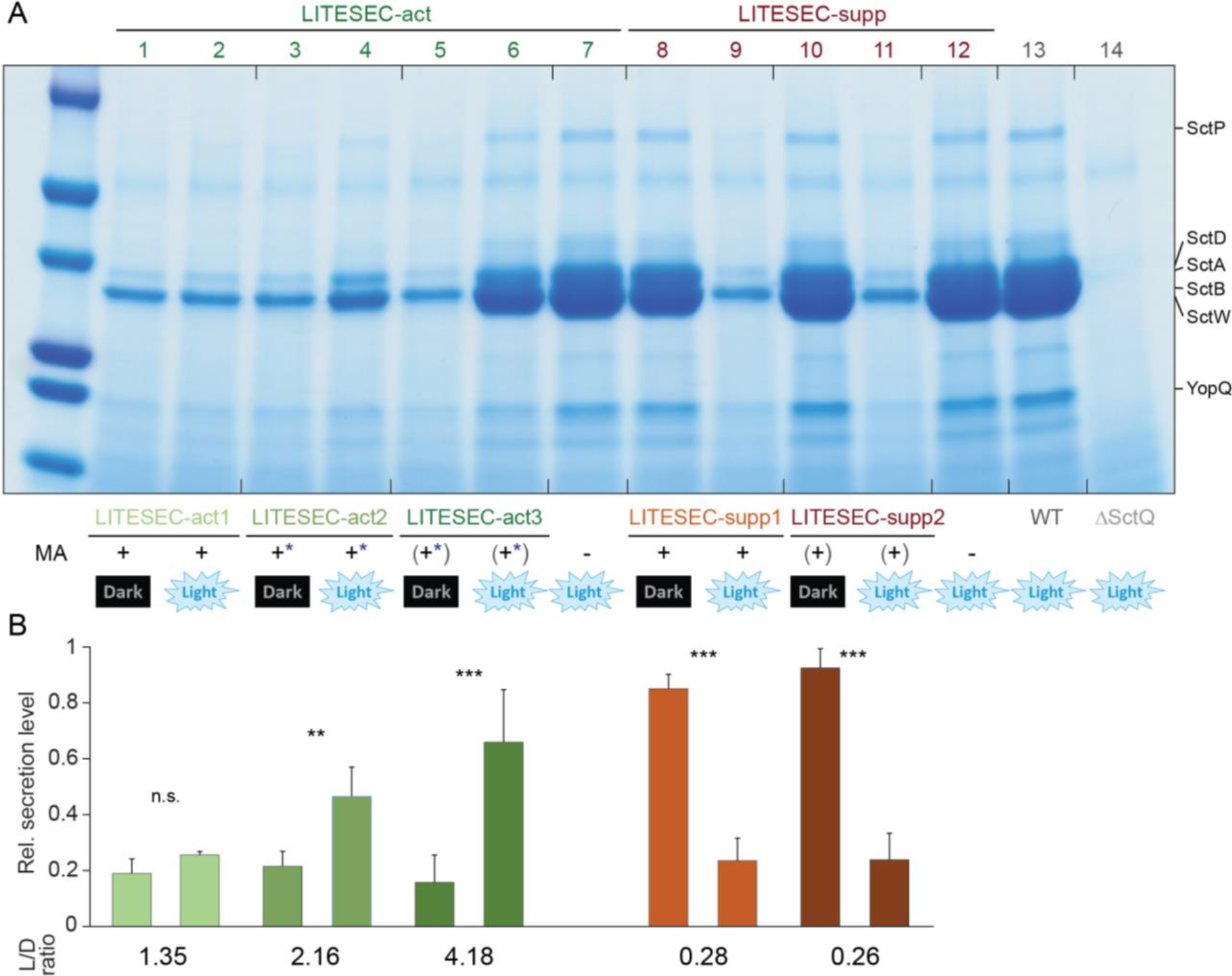
Secretion efficiency and light responsiveness in different versions of the LITESEC strains.

(**A**) *In vitro* secretion assay showing light-dependent export of native T3SS substrates (indicated on the right) in various variants of the LITESEC-act strains (lanes 1-7) and LITESEC-supp strains (lanes 8-12), as indicated below. Proteins secreted by 3^*^10^9^ bacteria during a 180 min incubation period were precipitated and analyzed by SDS-PAGE. MA, expression level of membrane anchor; +, high expression level; (+), low expression level; −, no expression. ^*^, V416L anchor mutant. (**B**) Quantification of secretion efficiency and light/dark secretion ratio (L/D ratio) for the different LITESEC strains and illuminations indicated above (as in (A)). Secretion efficiency was determined by gel densitometry for the YopB/LcrV/YopD/YopN bands and normalized for the secretion efficiency in wild-type strains (lane 13 in (A)), n=3−7, error bars display standard deviation. ^**^, p<0.01; ^***^, p<0.001 in a two-tailed homoscedastic t-test; n.s., difference not statistically significant.

### Light-dependent activation of the T3SS depends on the anchor/bait ratio

To assess whether the changed secretion efficiencies are indeed due to the lower expression of the anchor proteins in the new strains, we determined the expression levels by immunoblot. As expected, the anchor proteins expressed from the pBAD plasmids in the LITESEC-act2/-supp1 strains show a higher expression level than the anchor proteins expressed from the pACYC184 plasmid in the LITESEC-act3/-supp2 strains (Suppl. Fig. 4). To more thoroughly explore the connection between the anchor/bait expression ratio and the responsiveness of the T3SS to illumination, we compared the secretion levels under light and dark conditions for different expression levels of the anchor in the LITESEC-act2 system. The results show that indeed, the light responsiveness of the system (the difference between secretion levels under light and dark conditions) was optimal for intermediate anchor expression levels (Fig. 5A-C), corresponding to anchor/bait ratios of about one to two (Fig. 5D and Suppl. Methods)

**Fig. 5:**
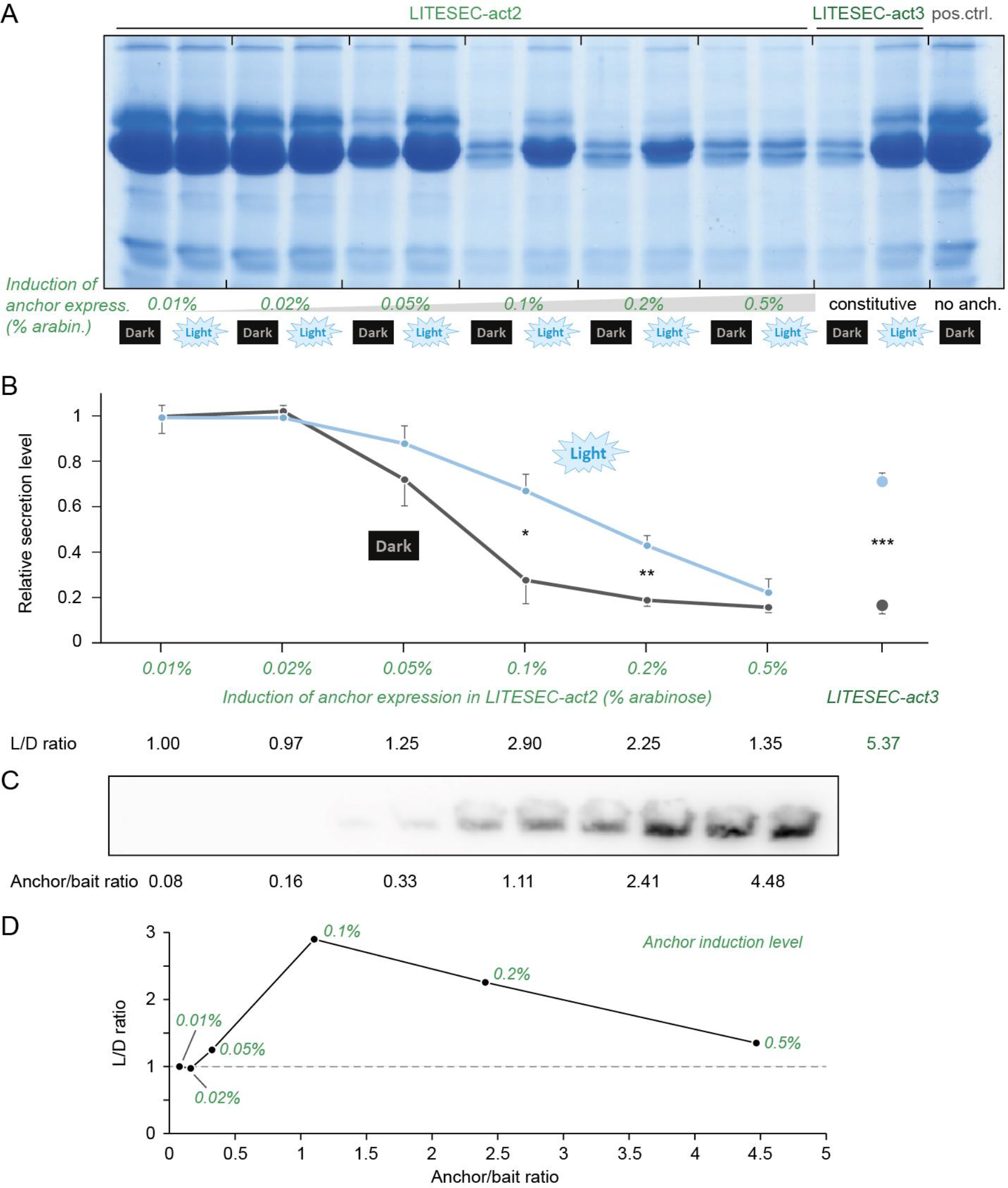
The expression ratio of anchor and bait protein dictates the function and light responsiveness of protein secretion in LITESEC-act2.

(**A**) *In vitro* secretion assay showing light-dependent export of native T3SS substrates in the LITESEC-act2 strain at different induction levels of anchor expression. (**B**) Quantification of secretion efficiency and light/dark secretion ratio (L/D ratio) for the different expression levels indicated above (as in (A)). ^*^/^**^/^***^, p<0.05/0.01/0.001 in a two-tailed homoscedastic t-test. (**C**) Western blot anti-FLAG of total cellular protein of 2^*^10^9^ bacteria in the indicated strains. Below, resulting anchor/bait ratio (see Suppl. Methods for details). (**D**) Correlation between light/dark secretion ratio (L/D ratio) and anchor/bait ratio. Labels indicate anchor induction levels (arabinose concentrations for LITESEC-act2); the grey dashed line denotes an L/D ratio of 1, indicating non-light-regulated secretion; n=3−4 for all experiments.

### The export of heterologous substrates by the T3SS can be controlled by light

The T3SS-dependent export of heterologous cargo has been shown and applied for many purposes in earlier studies^10,14,20^. To confirm that we can control the export of heterologous proteins in the LITESEC strains, we combined the LITESEC-act3 and -supp2 systems with a plasmid expressing a heterologous cargo protein, the luciferase NanoLuc, fused to a short N-terminal secretion signal, YopE_1-53_^16,40,41^, and a C-terminal FLAG tag for detection. The cargo protein was exclusively exported in light conditions by the LITESEC-act3 strain, and exclusively in the dark by the LITESEC-supp2 strain, whereas export was light-independent in a wild-type strain (Fig. 6).

**Fig. 6:**
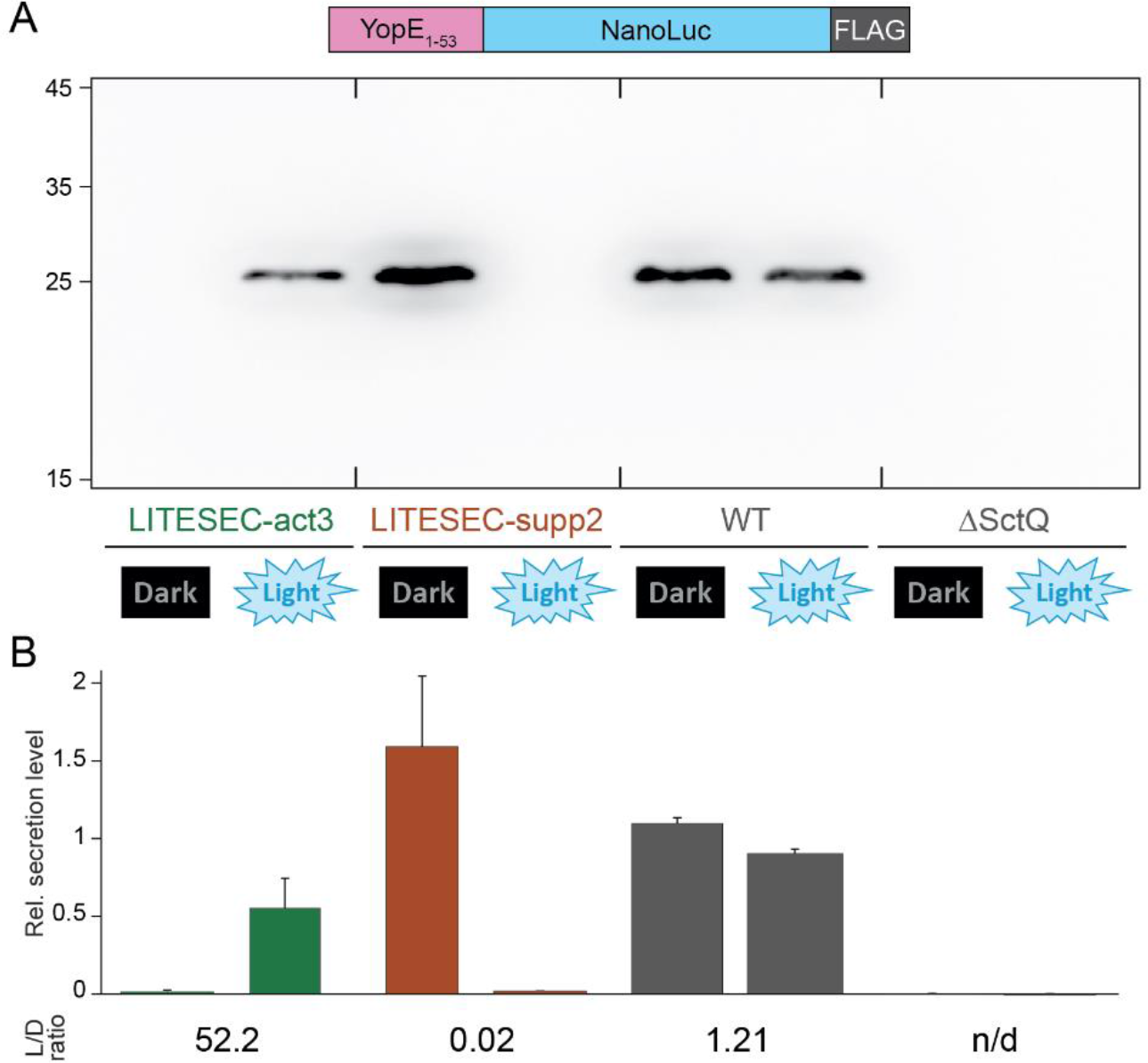
Heterologous cargo can be exported in a light-dependent manner.

*In vitro* secretion assay showing light-dependent export of YopE_1-53_-NanoLuc-FLAG (see scheme on top; exp. size, 28.7 kDa), in the indicated strains. Western blot using anti-FLAG antibodies. Left side, molecular weight in kDa. (**B**) Quantification of light-dependent YopE_1-53_-NanoLuc-FLAG export by densitometric analysis of Western blots, normalized by average secretion of the wild-type control (WT), n=3, error bars display the standard error of the mean. L/D ratio, ratio of secretion under light and dark conditions.

### Kinetics of light-induced T3SS activation and inactivation

How efficiently can the LITESEC system be inactivated, and what are the kinetics of light-induced T3SS activation and deactivation? Protein secretion for the LITESEC-act3 and -supp2 strains was analyzed for bacteria incubated consecutively for 60 min under inactivating conditions (dark for LITESEC-act3, light for LITESEC-supp2), 60 min under activating conditions, and another 60 min under inactivating conditions. After each incubation period, the culture medium was replaced, and a sample was tested for secretion by SDS-PAGE ad Western Blot. Secretion of the heterologous export substrate YopE_1-53_-NanoLuc-FLAG in LITESEC-act3 was specifically induced inlight conditions, and efficiently suppressed in the dark, whereas LITESEC-supp2 displayed the opposite behavior (Fig. 7). The WT strain continuously secreted proteins irrespective of the illumination. These results show that the activity of the LITESEC systems can be efficiently toggled. To more precisely determine the activation and deactivation kinetics, we used a sensitive bioluminescence-based luciferase assay^42^ to quantify the export of the reporter protein YopE_1-53_-NanoLuc-FLAG in the different LITESEC strains under changing illumination. In the LITESEC-supp2 strain, secretion of the heterologous substrate dropped to background levels within four to eight minutes after the start of blue light illumination, and recovered within the first four minutes after shifting the bacteria to dark conditions again. The LITESEC-act3 strain showed an increase of secretion activity over 20 minutes in light conditions, and required 12-16 minutes to shut down secretion in the dark (Suppl. Fig. 5).

**Fig. 7:**
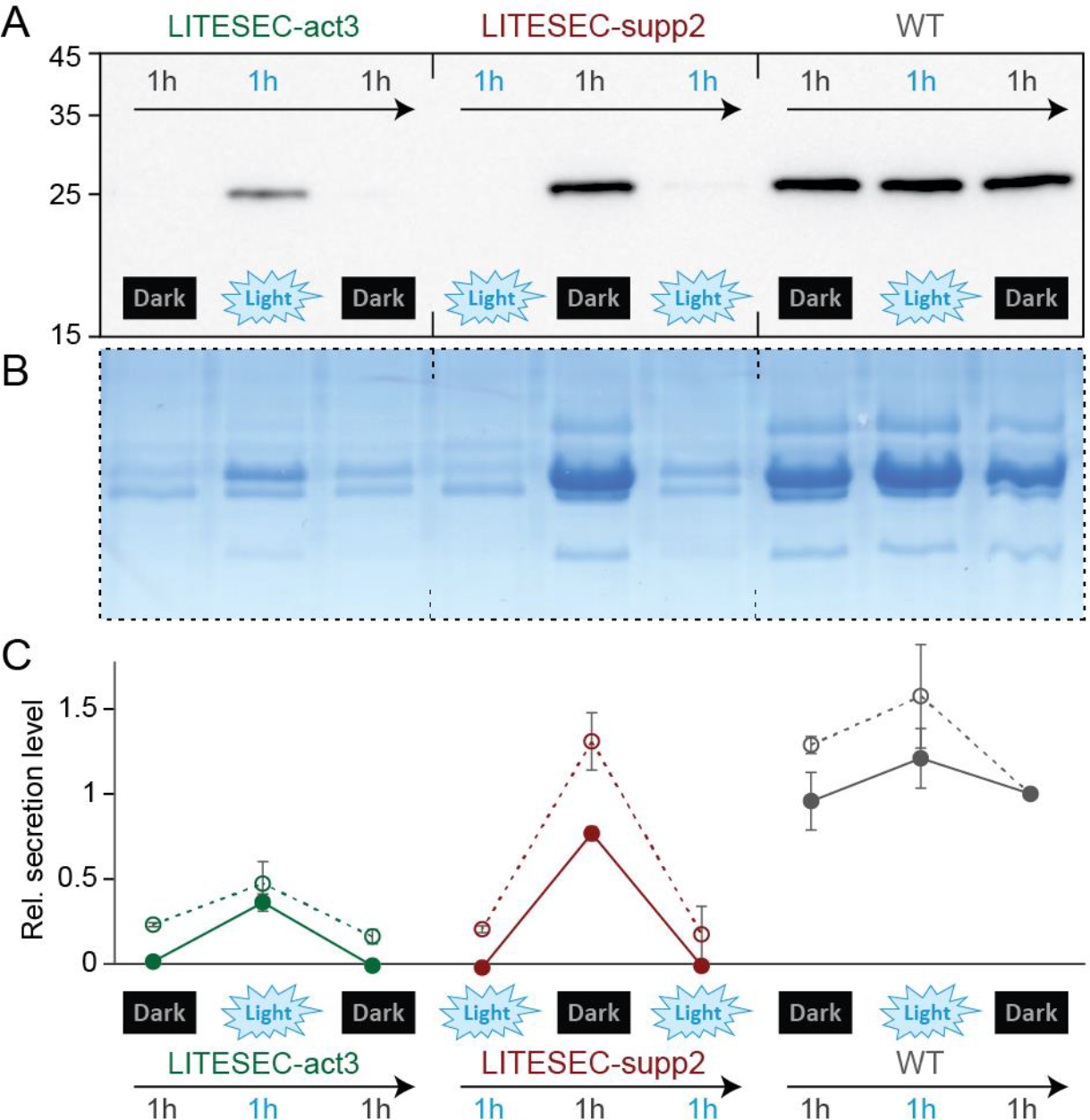
Secretion of effector proteins can be controlled by light over time.

(**A/B**) Export of the heterologous substrate YopE_1-53_-NanoLuc-FLAG (A) and native T3SS substrates (B) in the indicated strains. Secretion-competent bacteria were subsequently incubated under inactivating, activating and inactivating light conditions for 60 min each, as indicated. (**C**) Quantification of the relative export efficiency (normalized to the wild-type level in the third incubation period) of the strains and conditions shown in (A) and for export of YopE_1-53_-NanoLuc-FLAG (filled circles, continuous line) and endogenous T3SS translocator proteins (empty circles, dashes line); n=3, error bars denote standard deviation.

### Light-induced protein translocation into eukaryotic host cells

Having found that secretion of heterologous T3SS substrates can be tightly controlled by the LITESEC system, we wanted to employ the LITESEC-act system to control the injection of a cargo protein, YopE_1-53_-β-lactamase, into eukaryotic host cells upon illumination. Translocation of β-lactamase can be visualized by the cleavage of a Förster resonance energy transfer (FRET) reporter substrate, CCF2, within host cells^43,44^, which results in a green to blue shift in the emission wavelength. As expected, a wild-type strain translocated the YopE_1-53_-β-lactamase reporter substrate into a high fraction of host cells irrespective of the illumination. The negative control, the same strain expressing the β-lactamase reporter without a secretion signal, displayed a significantly lower rate of blue fluorescence (Fig. 8A), showing that translocation was T3SS-dependent. The LITESEC-act3 strain translocated the transporter in a light-dependent manner, leading to a significantly higher fraction of translocation-positive host cells in light than in dark conditions (close to the positive and negative controls, respectively; Fig. 8A). In contrast, the LITESEC-supp2 strain showed the opposite behavior (Fig. 8). Taken together, these results confirm that translocation of heterologous proteins into eukaryotic host cells by the T3SS can be controlled by external light.

**Fig. 8:**
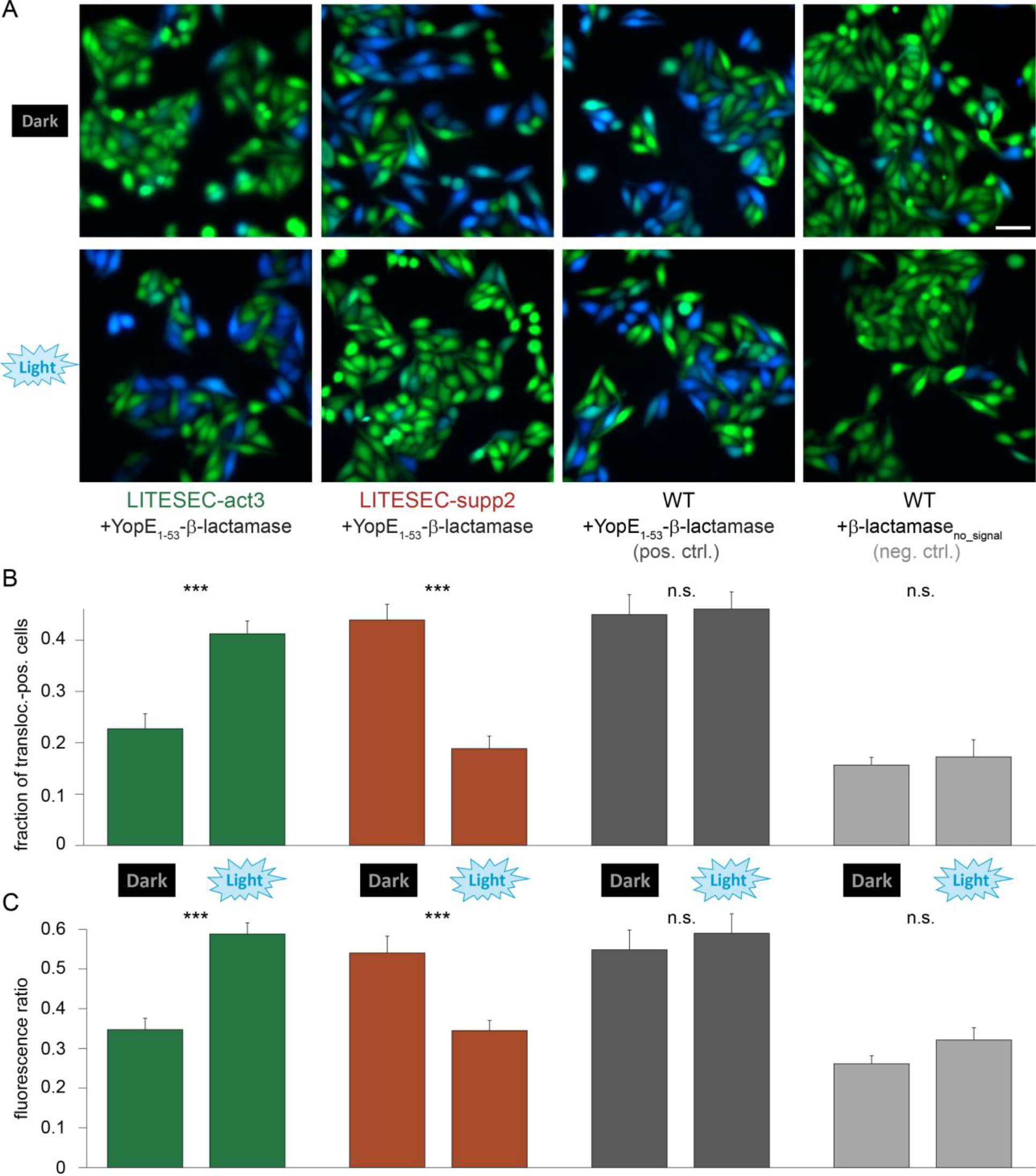
Light-dependent translocation of heterologous cargo into eukaryotic host cells.

(**A**) Fluorescence micrographs depicting cultured HEp-2 cells that have been incubated with the indicated LITESEC strains expressing either a heterologous T3SS substrate, YopE_1-53_-β-lactamase, or β-lactamase without a secretion signal as a negative control. Translocation of β-lactamase is detected by cleavage of the intracellular β-lactamase substrate CCF2 (leading to loss of FRET, and a transition from green to blue fluorescence emission). Scale bar, 50 μm. (**B**) Fraction of β-lactamase-positive HEp-2 cells (blue fluorescence). (**C**) Quantification of the fluorescence ratio of CCF2 donor fluorescence (indicative of β-lactamase translocation) and FRET fluorescence for (A). For panels B-C, 2226-2694 cells from 25-28 fields of view from 3 independent experiments were analyzed per strain and condition for the LITESEC strains (671-995 cells from 8-10 fields of view from 3 independent experiments for the controls). Error bars display the standard error of the mean amongst fields of view. ^***^, p<0.001 in a two-tailed homoscedastic t-test; n.s., difference not statistically significant.

## Discussion

(See Supplementary Information for a more detailed discussion of activation dynamics and applications of the LITESEC system.)

### Controlling protein secretion and translocation by the T3SS with light

To overcome the lack of specificity of T3SS-dependent protein secretion and translocation into eukaryotic cells, we aimed to control T3SS activity by external light. Our solution exploits the recently uncovered dynamic exchange of various essential T3SS components between an injectisome-bound state and a freely diffusing cytosolic state^25,31^ to control T3SS-dependent protein secretion by protein sequestration. SctQ, an essential and dynamic cytosolic component of the T3SS^31^, was genetically fused to one interaction domain of two optogenetic sequestration systems, the iLID and LOVTRAP systems^35,36,45^, while the membrane-bound interaction domain was co-expressed *in trans*. The two versions of the resulting LITESEC-T3SS system (**L**ight-**i**nduced **s**ecretion of **e**ffectors through **s**equestration of **e**ndogenous **c**omponents of the **T3SS**) can be applied in opposite directions: in the LITESEC-supp system, protein export is suppressed by blue light illumination; the LITESEC-act system allows to activate secretion by blue light.

The LITESEC-supp1 system, which is based on the iLID optogenetic interaction switch^34^ (Table 1), showed a significant reaction to light (light/dark secretion ratio of 0.28; 24% vs. 85% of wild-type secretion under light and dark conditions, respectively; Fig. 3). Expression of the membrane anchor from a constitutively active promoter on a low copy plasmid, pACYC184 (LITESEC-supp2) retained the light/dark secretion ratio (L/D ratio of 0.26; 24% vs. 93% WT secretion; Fig. 4), with the additional advantage that expression of the membrane anchor is constitutive.

For many applications, activation of T3SS protein export upon illumination is preferable. The LITESEC-act1 system, which is based on the LOV optogenetic interaction switch^35^, only achieved weak activation of T3SS secretion upon illumination (Fig. 4). LITESEC-act2, which uses the V416L mutation in the anchor protein^46^ to decrease the affinity between anchor and bait, could be activated by light more efficiently. Even more strikingly, LITESEC-act3, featuring a reduced expression level of the V416L variant of the membrane anchor, led to a strong activation of T3SS protein secretion upon illumination, while retaining the tight suppression of secretion in the dark (L/D ratio of 4.2; 66% vs. 16%; Fig. 4).

Notably, the export of heterologous cargo was entirely light-dependent (no visible export under inactive conditions; Fig. 6). In contrast, endogenous T3SS translocator proteins were still secreted to a basal level under inactivating light conditions, even in the most tightly controlled strains (LITESEC-act3/-supp2; Fig. 4). This might indicate that the export of heterologous cargo is regulated differently from the export of the endogenous translocators, which for example also involves protein-specific chaperones. While this hypothesis remains to be rigorously tested, it highlights that beyond their application, LITESEC and similar optogenetic approaches can help to better understand the underlying biological systems.

To explore the influence of the anchor/bait expression ratio on light control of the T3SS in more detail, we measured the light-dependent activation of the LITESEC-act2 system at different expression levels of the anchor protein. The results indicate that anchor/bait ratios of around one to two allowed an optimal response to blue light for the LITESEC-act system. Higher ratios retain partial membrane sequestration under light conditions and subsequently impair T3SS activity in the activated stage; conversely, low ratios lead to incomplete sequestration and measurable T3SS activity under non-activating conditions (Fig. 5). Due to possible variations in transfer efficiency and the indirect nature of the anchor/bait ratio determination, this value might not be precise; however, our data strongly suggest a relatively tight “sweet spot” in the expression ratio of the two interacting proteins, which may be key for the successful optogenetic control of bacterial processes. This is in sharp contrast to the eukaryotic application of the LOVTRAP interaction switches where much higher anchor/bait concentrations where shown to be optimal^35^. We therefore propose that optimization of the anchor/bait expression ratio represents an important step in the design of optogenetically controlled processes in prokaryotes.

### Factors for controlling prokaryotic processes by optogenetic interaction switches

The successful development and application of the LITESEC system highlights some key features for the control of prokaryotic processes by optogenetic interaction switches. The target protein (in our case the essential T3SS component SctQ) (i) has to be functional as a fusion protein to an optogenetic interaction domain, (ii) must be present in the cytosol at least temporarily to allow sequestration to occur, and (iii) must not be functional when tethered to the membrane anchor protein. To fulfil the last criterion, the target protein may feature a) a specific place of action (such as the injectisome for SctQ in the present case), or b) a specific interaction interface that is rendered inaccessible by the interaction with the anchor. In eukaryotic systems, proteins have been sequestered to various structures including the plasma membrane or mitochondria. The simpler cellular organization of bacteria makes the inner membrane an obvious target for protein sequestration, to which interaction domains can be easily targeted to by the addition of N-terminal TMHs. While the nature of the TMH is likely to be secondary for the success of the application, the extended TatA TMH and a short glycine-rich linker worked well for our approach. Crucially, we found that the expression ratio between anchor and bait proteins is a key determinant for the success of LITESEC and, most, likely, similar approaches to control bacterial processes by light.

### Light-controlled protein translocation into host cells

The T3SS is a very promising tool for protein delivery into eukaryotic cells, both in cell culture and in healthcare^10,14,20^. However, the T3SS indiscriminately injects cargo proteins into contacting host cells. Lack of targetability is therefore a main obstacle in the further development and application of this method^20,21^. Previous methods to control the activity of the T3SS relied on controlled expression of one or all components of the injectisome. For example, Song and colleagues expressed all components of the *Salmonella* SPI-1 T3SS from two inducible promoters in a clean expression system^47^, and Schulte *et al*. expressed the T3SS genes from a TetA promoter, which additionally allows the intracellular induction of the T3SS^48^. Besides the difficulty to specifically induce secretion in defined places *in situ*, the main drawback of these methods is the slow response (induction of expression and assembly of the T3SS take more than 60 min^28,47,48^). In addition, in these systems, the T3SS remains active as long as the induced protein(s) are still present, which leads to a higher risk of translocation into non-target cells.

By using light to specifically activate the modified T3SS in bacteria, we have addressed this issue. The LITESEC system allows to deliver proteins into host cells at a specific time and place. The system gives complete control over the secretion of heterologous T3SS cargo into the supernatant, either by providing illumination (LITESEC-act), or stopping the light exposure (LITESEC-supp). Importantly, secretion by the LITESEC-act system is temporary, and stopped within minutes after the end of illumination with blue light, thereby further reducing unspecific activation.

The LITESEC system presented in this work uses light-controlled sequestration of an essential dynamic T3SS component to precisely regulate the activity of the T3SS. This approach provides a new method for highly time- and space-resolved protein secretion and delivery into eukaryotic cells.

Supporting information, including material and methods, can be found in the supplementary file.

## Acknowledgements

The authors would like to thank Prof. Andreas Gahlmann, University of Virginia, for important contributions and discussions during the inception of this study; Dr. Felicity Alcock, University of Oxford, for suggestions for the design of the transmembrane anchor; Dr. Seraphine Wegner, Max-Planck-Institute for Polymer Research, Mainz for helpful discussions about optogenetic interaction switches; Prof. Petra Dersch, University of Münster, Germany, for the gift of HEp-2 cells; as well as Prof. Lars-Oliver Essen, University of Marburg, for input and support on the characterization of the optogenetic setup.

## Competing interests

A. Diepold and F. Lindner submitted a European patent application on the presented method. The authors declare no competing financial interests.

## Footnotes

§ In this manuscript, T3SS refers to the virulence-associated T3SS. The common „Sct‟ nomenclature^49^ is used for T3SS components, see ref. ^39^ for species-specific names.

