## Supplementary Information for "LITESEC-T3SS - Light-controlled protein delivery into eukaryotic cells with high spatial and temporal resolution"

### Supplementary Methods

Plasmids and strains used in this study are listed in Suppl. Tables 2 and 3, respectively.

*Cultivation of bacteria*

*Y. enterocolitica* strains were cultivated in rich BHI (Brain Heart Infusion Broth) medium (3.7% w/v), complemented with nalidixic acid (35 µg/ml) and 2,6-diaminopimelic acid (DAP, 60 µg/ml) (cultivation medium). *E. coli* strains were cultivated in LB (Lurea Broth) medium (tryptone (10% w/v), yeast extract (5% w/v), NaCl (10% w/v)). Where required, the medium was supplemented with ampicillin or carbenicillin (200 µg/ml), chloramphenicol (25 µg/ml), or streptomycin (50 µg/ml). For overnight cultures, 2-5 ml of cultivation media with corresponding antibiotics were inoculated and cultivated over night at 28°C (*Y. enterocolitica*) or 37°C (*E. coli*) in a shaking incubator.

*T3SS in vitro secretion assay*

Day cultures were inoculated from stationary overnight cultures (1:50 dilution for non-secreting conditions, 1:41.67 for secreting conditions) in cultivation medium additionally complemented with $MgCl_2$ (20 mM), glycerol (0.4% w/v), and corresponding antibiotics. $CaCl_2$ (5 mM) or EGTA (5 mM) were added for non-secreting and secreting conditions, respectively. The cultures were cultivated for 90 min at 28°C and then shifted to a 37°C water bath and inoculated for 2-3 h (if the strain contained an inducible plasmid, the plasmid was induced with 0.2% w/v L-Arabinose before shifting to 37°C).

*Fluorescence microscopy*

For fluorescence microscopy, strains were cultivated as described above under non-secreting conditions. 2 ml of culture then was harvested for 4 min at 2,400 *g* and the cell pellet was resuspended in 400 µl of minimal medium (HEPES (100 mM), $(NH_4)_2SO_4$ (5 mM), NaCl (100 mM), sodium glutamate (20 mM), $MgCl_2$ (10 mM), $K_2SO_4$ (5 mM), casamino acids (0.5% w/v)) including DAP (60 µg/ml). From this culture, 2 µl were spotted onto agar slides (1.5% w/v agarose in minimal media and topped with a circular cover slip (25 mm ⌀). Samples were analyzed on an inverse fluorescence microscope (Deltavision Elite). For pulse activation of the optogenetic interaction switches, 0.1 s of GFP excitation light (~ 480 nm, light intensity ~ 2.5 mW/cm$^2$) was applied. Unless stated differently, exposure times were 500 ms for mCherry fluorescence, using a mCherry filter set, and 200 ms for GFP fluorescence, using a GFP filter set. Per image, a z stack containing 7 to 15 frames per wavelength with a spacing of 150 nm was acquired.

*Optogenetic cell cultivation*

For optogenetic experiments, the strains for secretion assays or Western blots (to determine the amount of secreted proteins) were cultivated under secreting conditions as described before. At the indicated time points after induction of the system by a temperature shift to 37°, the cultures were cultivated at 37°C for 1 - 3 h in an optogenetic experimental setup (Suppl. Fig. 7) under blue light or dark conditions (light intensity at culture location at a wavelength of 488 nm was ~ 1 mW/cm$^2$), and further processed as described.

*Infection assay*

HEp-2 cells were maintained in Roswell Park Memorial Institute (RPMI) 1640 medium (Gibco) supplemented with 7.5% newborn calf serum (NCS, Sigma-Aldrich ) in 5% $CO_2$ at 37°C. HEp-2 cells were seeded into Nuncion Delta Surface 96- flat well plates (Thermo Scientific) at a cell density of 2.0x10$^4$ cells/well. Prior to infection, 5 mM DAP was added to medium of the seeded HEp-2 cells.

The infection assay was adapted from ref. 1. 200 µl of bacterial overnight culture was inoculated in BHI supplemented with DAP (60 µg/ml), $MgCl_2$ (20 mM), and glycerol (0.4% w/v). Expression of the cargo protein from the pBAD plasmid was induced with 0.2% arabinose (w/v), unless stated differently. The cultures were incubated for 90 min at 37°C under activating conditions (dark for LITESEC-supp / light for LITESEC-act) to induce T3SS formation. After incubation, cultures were centrifuged for 4 min at 4,500 *g* and 4°C. Cells were resuspended in ice-cold PBS containing 5 mM DAP at a density of approximately 2.5x10$^8$ cfu/ml. Bacteria were incubated on ice in “off” conditions (light for LITESEC-supp / dark for LITESEC-act) for 15 min, then added to a semi-confluent layer of HEp2-cells at an MOI of approximately 140, and incubated under blue light or dark conditions for 60 min at 37°C in 5% $CO_2$. Following incubation, the cell culture medium was removed and 100 µl of working solution were added (1:3 dilution RPMI 1640 medium without phenol red (Gibco) in PBS (Gibco) with 25 mM probenecid acid (Alfa Aesar) dissolved in cell culture grade DMSO (Chem Cruz)). 20 µl of CCF2-AM were added (0.12 µl solution A, 1.2 µl solution B and 18.68 µl solution C (solutions A, B and C provided from Invitrogen CCF2-AM loading kit)). After 5 min of incubation, the working solution and CCF2-AM were removed and 100 µl of fresh working solution was added. Plates were then incubated at 37°C in 5% $CO_2$ for 10 min. Next, cells were fixed by addition of 100 µl of ice-cold 1% para-formaldehyde (PFA) (w/v) and incubation on ice for 10 min. As a last step, the PFA solution was replaced by PBS. Translocation of YopE$_{1\text{-}53}$-β-lactamase was detected by comparing the fluorescence emission at 525/48 nm (FRET-based emission of uncleaved CCF2) vs. 435/48 nm (emission of cleaved

CCF2, equivalent to substrate translocation), both at an excitation at 390/18 nm. Both channels were background-corrected. The fluorescence of HEp-2 cells was manually classified by blinded observers.

*Luciferase assay*

20 ml day cultures were inoculated from stationary overnight cultures (1:50 dilution) in non-secreting cultivation medium, as described above, and incubated for 90 min at 28°. Subsequently, expression of the $YopE_{1\text{-}53}$-NanoLuc-Flag cargo protein from the pBAD plasmid was induced with 0.2% arabinose (w/v). The cultures were incubated for 120 min at 37°C under activating conditions (dark for LITESEC-supp / light for LITESEC-act) to induce T3SS formation. After that, strains were incubated for 10 min under dark conditions, and 5 mM EGTA was added to start secretion. Bacteria were then incubated for 20 min each under activating, inactivating, and activating conditions. Samples were removed from the cultures immediately before EGTA addition and every 4 min afterwards. In the samples, 10 mM $CaCl_2$ was added to stop secretion. Bacteria were harvested and the supernatant was used for the enzymatic assay. The enzymatic Nanoluc detection assay was performed according to manufacturer instructions. Briefly, 5 µl supernatant was mixed with 25 µl $H_2O$ and 30 µl of Nanoluc-detection reagent (Nano-Glo Luciferase Assay Substrate – Nano-Glo Luciferase Assay Buffer 1:50 – Nano-Glo Luciferase assay kit (PROMEGA Corporation, Madison)). Bioluminescence was detected via Elisa Plate Reader Infinite M20 Pro (BioTek Instruments, Vermont), smallest field of view, large binning and an aquisition time of 1,000 ms.

*Calculation of anchor/bait ratio*

To calculate the ratio of the anchor and bait proteins, the band intensities of endogenously expressed Zdk1-SctQ-mCherry, and the membrane anchor TMH-FLAG-LOV2-mCherry, coexpressed *in trans* (Suppl. Fig. 4B), were quantified in an immunoblot anti-mCherry. To calculate the anchor/bait expresssion ratio in the LITESEC-act2 strain at an induction level of 0.2% arabinose, this result was corrected for the expression ratio of TMH-FLAG-$LOV2_{V416L}$ and TMH-FLAG-LOV2 (Suppl. Fig. 4A), determined in an anti-Flag immunoblot. All individual experiments were done in biological triplicates, the results are displayed in Suppl. Table 5. The resulting anchor/bait ratio for the LITESEC-act2 system induced with 0.2% arabinose was then used as a reference to determine the anchor/bait ratio at different arabinose induction levels by comparing the expression levels of TMH-FLAG-$LOV2_{V416L}$ under dark conditions at these different induction levels (Fig. 5C), and for the correlation of L/D activity ratios and anchor/bait ratios (Fig. 5D).

### Supplementary Discussion

*Kinetics of LITESEC activation and deactivation*

Both reaction time and recovery dynamics of the sequestration systems are crucial for their applicability to control the function of the T3SS. Fast reaction times to blue light increase the temporal precision of T3SS activation/deactivation, whereas the recovery times influence the duration of the effect on secretion after illumination. Very fast recovery means that the system has to be continuously illuminated for a sustained effect on secretion, while very slow recovery leads to long-term activation/deactivation that is difficult to revert, and renders handling of the cultures difficult due to possible long-term effects of illumination prior to the actual experiment. In time-course experiments we could show that in the iLID-based protein sequestration system, unbinding of the bait in the light state was almost immediate, and that recovery in the dark occurred within few minutes (Fig. 2), in line with data from eukaryotic systems[2]. In the resulting LITESEC-supp system, both activation and deactivation of type III secretion occur relatively quickly, within the first minutes (Fig. 7), which is in the range of the measured turnover of SctQ at the injectisome (half-time of about 70 s under secreting conditions[3]). This suggests that the release and rebinding of the bait protein occurs faster or in a similar time range, consistent with the microscopy results (Fig. 2). For the LITESEC-act system, we detected a slower activation and deactivation of protein secretion (Fig. 7). Nevertheless, induction of protein secretion by blue light occurs within minutes (Fig. 7). Importantly, in the absence of further illumination, protein secretion is stopped within minutes, which greatly limits unwanted unspecific activation. Long-term activation can be achieved by either constant low-intensity blue light illumination, or short light pulses every few minutes. Ambient laboratory light did not inhibit the LITESEC-supp2 strains, but lead to an intermediary activation of LITESEC-act3 (Suppl. Fig. 6).

*Current and possible future applications of the LITESEC system*

A main application of the LITESEC system is the temporally and spatially controlled translocation of proteins into cultured eukaryotic cells (Fig. 8). Cell cultures play an important role in research, development and, increasingly, healthcare. Often, specific proteins need to be expressed in all or a subset of the cultured cells at a given time point. At the moment, this is mainly achieved by inducing expression of the target protein within the host cells. This method requires prior transfection of the host cells with the target gene or time-consuming creation of stable transgenic cell lines. Induction of expression itself is relatively slow, and difficult to apply to a certain subset of cells. Our method allows to translocate proteins into unmodified host cells with high specificity. Bacteria that lack their native virulence effectors (such as the *Y. enterocolitica* strain used in this study), but express one or more cargo proteins with a short secretion signal, are brought into contact with host cells. The chosen subset of host cells are then subjected to dark or blue light conditions (which does not influence bacteria or host cells at the used intensity), which temporarily induces translocation of the cargo into the host cells within short time. An additional advantage of the LITESEC method is that it directly translocates proteins into the host cell, rather than inducing the transcription of mRNA, as is the case in the current inducible transfection systems. The amount of translocated protein can be regulated by the duration of illumination/darkness, and the multiplicity of infection (ratio of bacteria / host cells)[4].

A potential, relatively straightforward extension of our work would allow the specific protein delivery into diseased cells, such as cancer cells, within biological tissues. The T3SS has been used to treat cancer cells *in vitro*, e.g. by translocating angiogenic inhibitors[5], but again, the promiscuity of the T3SS and the resulting unspecific translocation at non-target sites represent a major obstacle in the further development of T3SS-based methods for clinical applications[6]. Most current approaches rely on localized injection of bacteria or the natural tropism of bacteria to tumorous tissue. However, bacteria applied with these methods are not restricted to the target tissue, and unspecific activation presents a problem, especially for potentially powerful applications such as the delivery of pro-apoptotic proteins. By using light to specifically activate the modified T3SS in bacteria at a site of choice, delivery of effector proteins could be temporarily induced at a specific time and place. This method would reduce unspecific activation and side effects, allowing a highly controlled targeting of host cells. Bacteria could be applied to the patient (exploiting the natural tropism of bacteria for tumor tissue to achieve local enrichment in the case of cancer), where injection of the effector protein would be triggered *in situ* with high spatial and temporal precision using light delivered with the help of endoscopes and minimally-invasive surgery techniques. As the blue light used to control the current LITESEC systems does not penetrate tissue efficiently, activation by red or far-red light would be advantageous. Several such red-light systems have been characterized[7–9]; however, all of these systems require cofactors not usually present in bacteria.

### Supplementary Figures

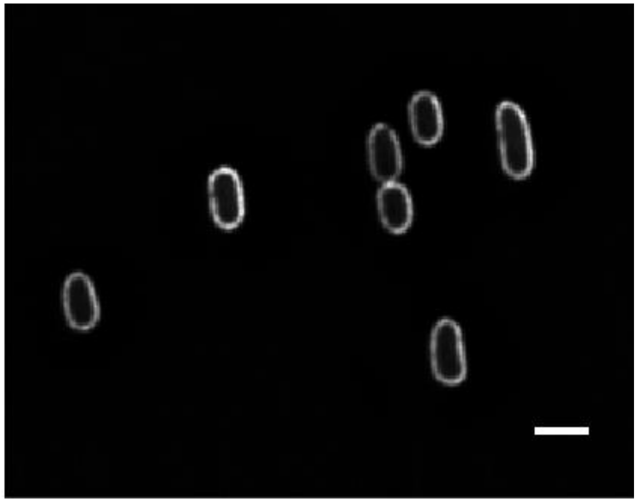

**Suppl. Fig. 1: Membrane localization of the LOV anchor protein in *Y. enterocolitica***

Localization of an mCherry-labeled version of the membrane anchor protein for the LOV-based membrane sequestration system, TMH-FLAG-mCherry-LOV, determined by widefield fluorescence microscopy. Scale bar, 2 µm.

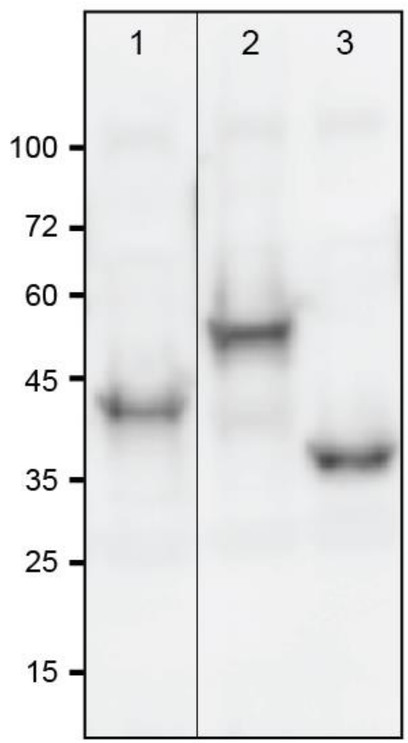


**Suppl. Fig. 2 – The used fusion proteins are stable.**

Western blot anti-SctQ of the used bait-SctQ fusion proteins, expressed from the native genetic locus on the virulence plasmid. Control strain, dHOPEMTasd (wild-type SctQ). Detected proteins and expected sizes: 1, Zdk1-SctQ, 40.8 kDa; 2, SspB_Nano-SctQ, 46.7 kDa; 3, WT SctQ, 34.4 kDa. The thin horizontal line denotes the omission of intermediate lanes on the same immunoblot.

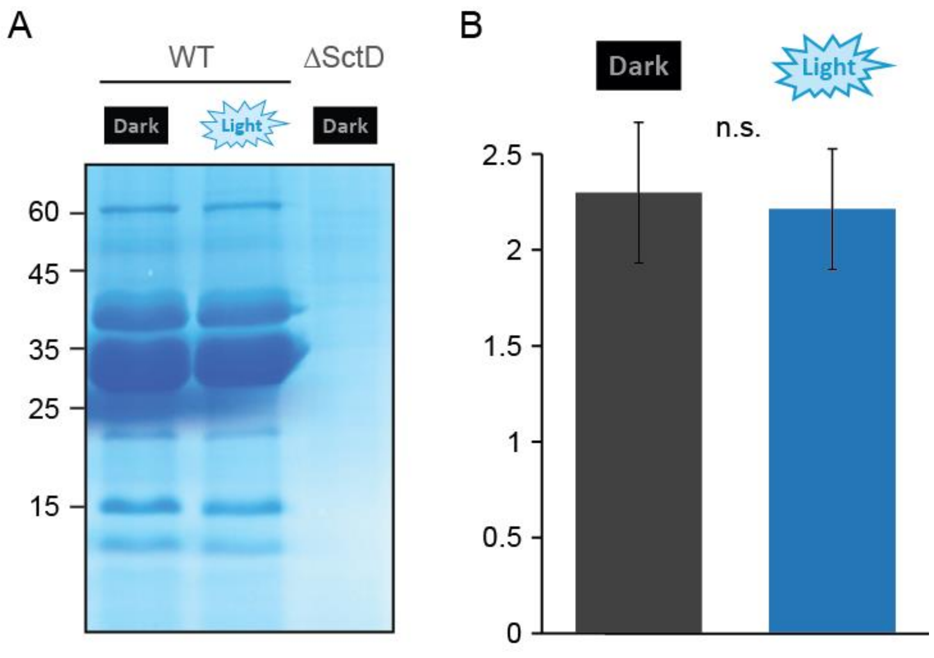


**Suppl. Fig. 3: Blue light illumination in the used intensity does not significantly influence growth, division, or T3SS activity of *Y. enterocolitica***

(**A**) *In vitro* secretion assay showing export of T3SS substrates in the indicated strains (WT, wild-type; ΔSctD, T3SS-negative control) in dark or light (constant low-level illumination of ~ 1 mW/cm$^2$ at $\lambda$=488 nm) conditions, as indicated. Proteins secreted by 3*10$^9$ bacteria during a 180 min incubation period were precipitated and analyzed by SDS-PAGE. (**B**) Average optical density at 600 nm of wild-type cultures in secreting conditions after 180 min in dark conditions (grey, left) or light conditions (blue, right) as used in the optogenetics experiments. n=3, n.s., no statistically significant difference (p=0.77 in a two-tailed homoscedastic t-test).

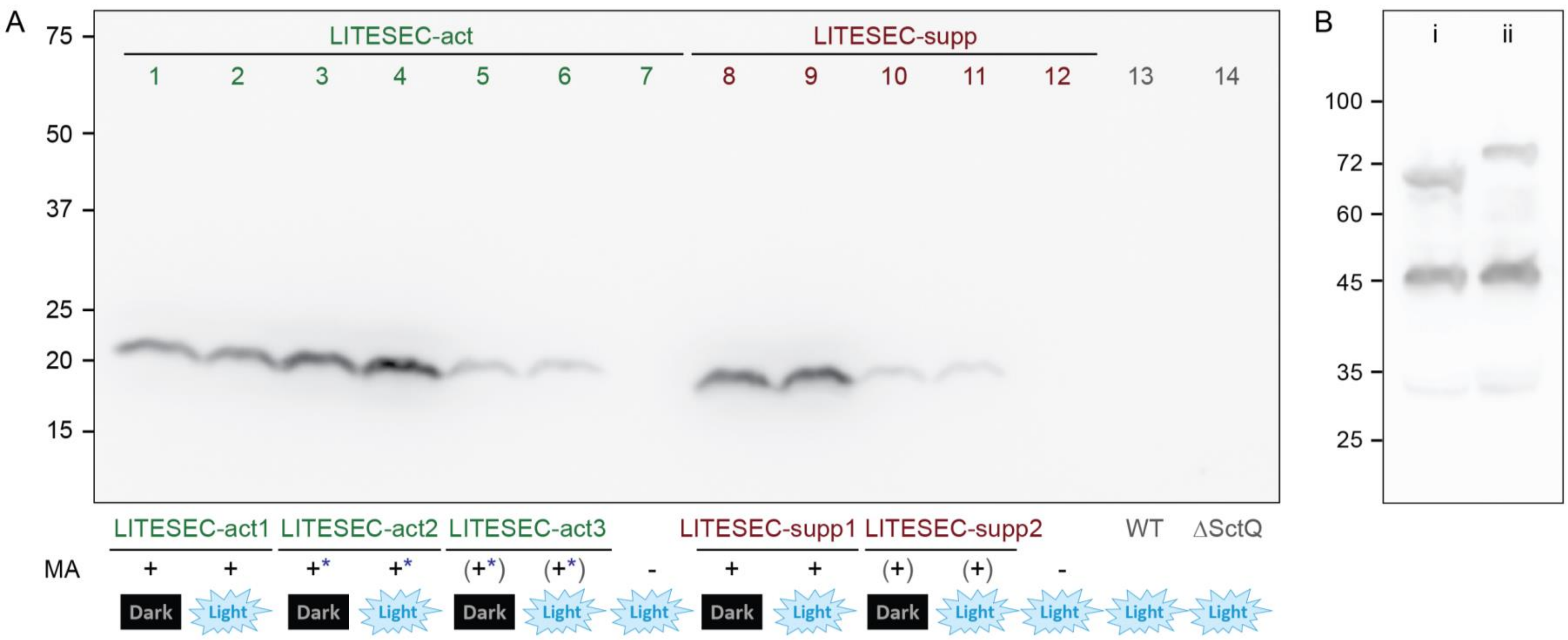


**Suppl. Fig. 4: Expression levels of membrane anchor proteins in the different LITESEC variant strains**

(**A**) Western blot anti-FLAG of total cellular protein from $2*10^9$ bacteria in the indicated strains (corresponding to Fig. 4). Left, molecular weight marker in kDa. Expected protein sizes: 20.9 kDa for LITESEC-act strains (TMH-FLAG-LOV2 / TMH-FLAG-LOV2$_{V416L}$), 21.6 kDa for LITESEC-supp strains (TMH-FLAG-iLID). (**B**) Western blot anti-mCherry of mCherry-labeled anchor and bait combinations of both LITESEC systems. Detected proteins and expected sizes: i, Zdk1-mCherry-SctQ and TMH-FLAG-mCherry-LOV2, 67,8 and 47,6 kDa; ii, SspB_Nano-mCherry-SctQ and TMH-FLAG-mCherry-iLID, 73,7 kDa and 48,7 kDa. Left, molecular weight marker in kDa. N=3.

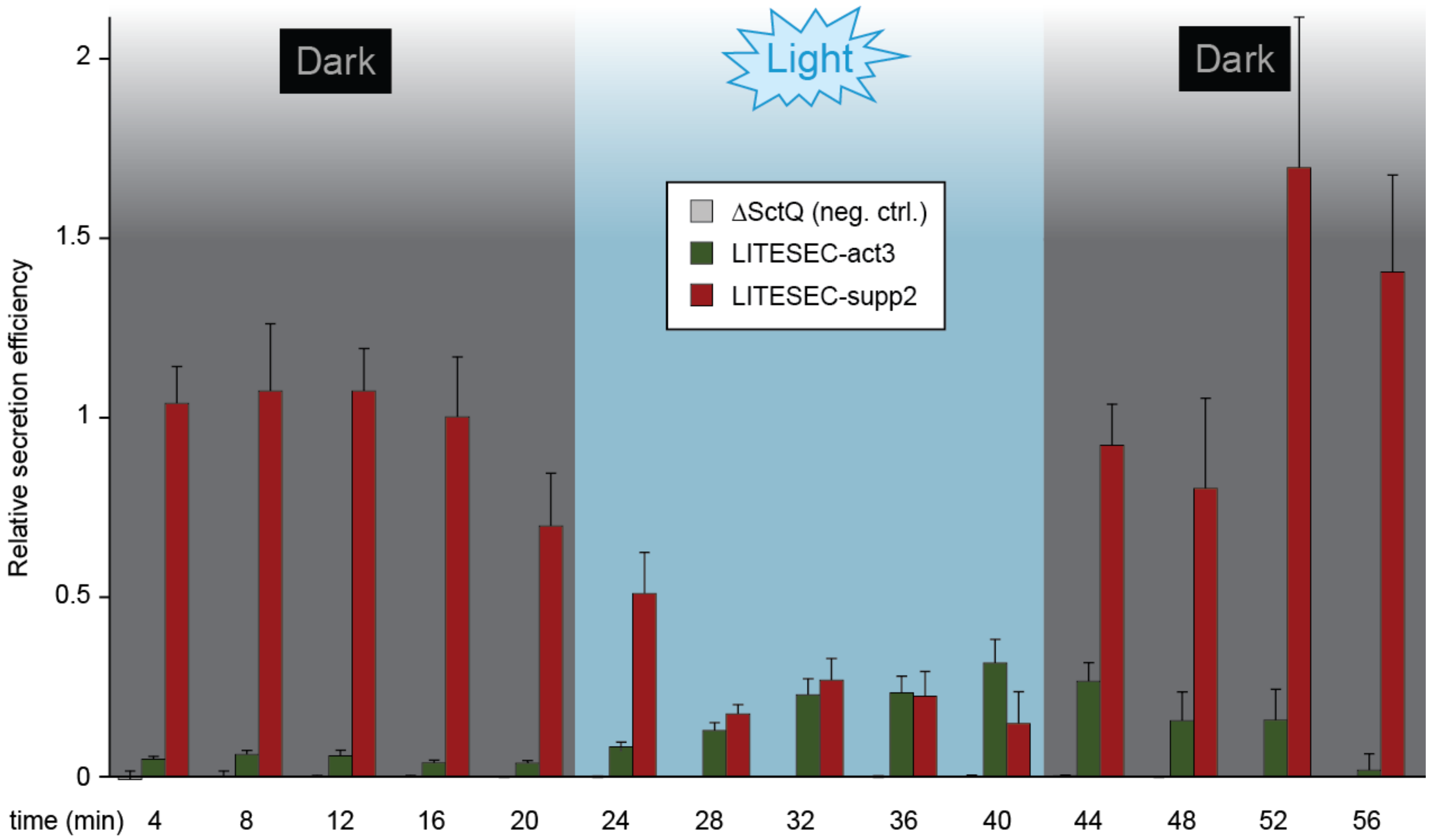


**Suppl. Fig. 5: Determination of switching kinetics in the LITESEC strains**

Secretion-competent bacteria were incubated under the indicated conditions. Every four minutes, a sample was removed, secretion was stopped by addition of 10 mM $CaCl_2$, and bacteria were harvested by centrifugation. Secretion of $YopE_{1\text{-}53}$-NanoLuc-FLAG was quantified by a Luciferase-based luminescence assay in a plate reader. The increase in secretion in comparison to the previous time point was determined and normalized for the secretion of a wild-type strain. N=5, error bars denote the standard error of the mean.

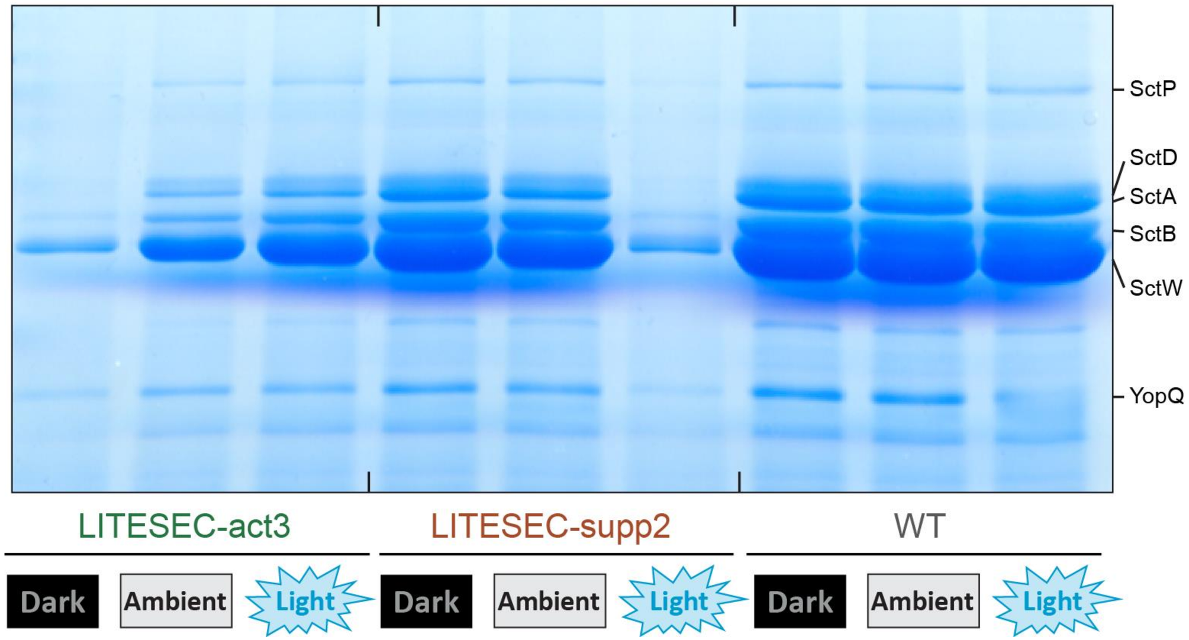


**Suppl. Fig. 6: Influence of ambient light on LITESEC activity**

*In vitro* secretion assay showing light-dependent export of native T3SS substrates (indicated on the right) in the listed strains, incubated under defined dark or light conditons (see material and methods for details), as well as ambient laboratory light. Proteins secreted by $3*10^9$ bacteria during a 180 min incubation period were precipitated and analyzed by SDS-PAGE.

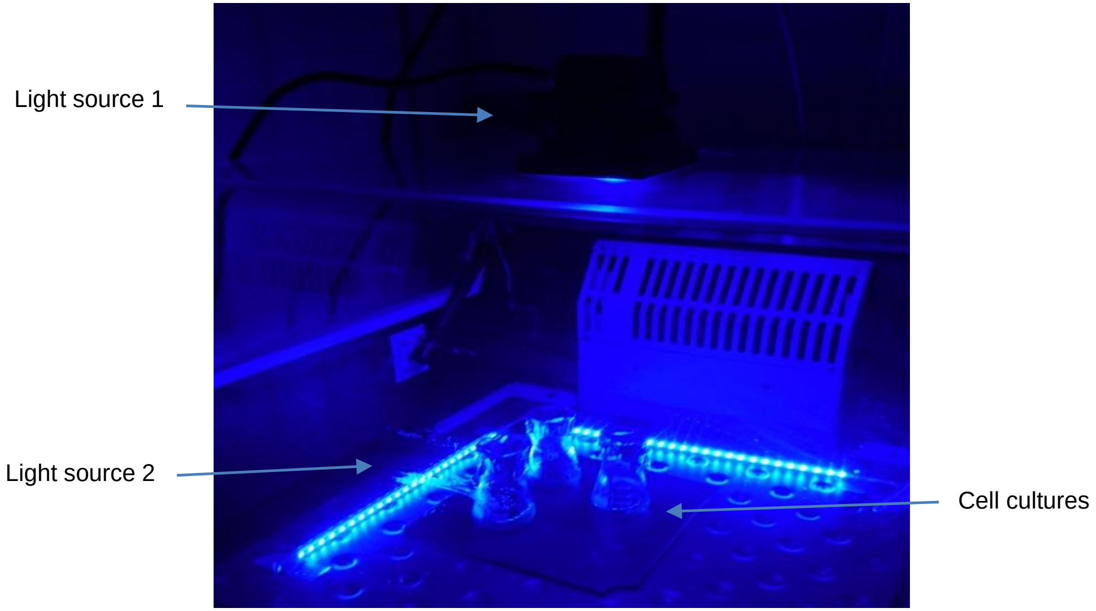


**Suppl. Fig. 7: Optogenetic experimental setup**

The optogenetic experimental setup consists of two blue light sources that were placed around the bacterial cultures. Light source 1 was a "globo lighting 10 W LED 9 V 34118S" – (Globo Lighting GmbH (St. Peter, A)), Light source 2 was a "Rolux LED-Leiste DF-7024-12 V 1.5 W" – (Rolux Leuchten GmbH (Weyhe, G)). Bacteria were cultivated at 37°C.

### Supplementary Tables

| Optogenetic system | Role of expressed protein | Plasmid name | Domains of expressed protein |
|---|---|---|---|
| **LOV** | *Anchor* | *pAD608* | *TMH-FLAG-LOV2* |
| | | *pFL100* | *TMH-FLAG-mCherry-LOV2* |
| | Bait | pFL104 | Zdk1-mCherry |
| **iLID** | *Anchor* | *pFL108* | *TMH-FLAG-iLID* |
| | | *pFL107* | *TMH-FLAG-mCherry-iLID* |
| | Bait | pFL109 | SspB_Nano-mCherry |

#### Suppl. Table 1: Optogenetic constructs for membrane sequestration assay

Constructs of the interaction partners used for the membrane sequestration assay, and their domains. TMH, extended TatA transmembrane helix (see material and methods for details).

| Name | Genotype | Strain background | Comments* / Reference |
|---|---|---|---|
| **dHOPEMTasd** | *pYV40 yopO*$_{Δ2\text{-}427}$ *yopE*$_{21}$ *yopH*$_{Δ1\text{-}352}$ *yopM*$_{23}$ *yopP*$_{23}$ *yopT*$_{135}$ *Δasd* | | Ref. 10 |
| **AD4324** | *mCherry-SctQ* | dHOPEMTasd | Ref. 3 |
| **AD4419** | *ΔSctQ* | dHOPEMTasd | Ref. 3 |
| **ADMT4521** | *mCherry-SctL* | dHOPEMTasd | Ref. 11 |
| **FL4002** | *Zdk1-mCherry-SctQ* | dHOPEMTasd | pFL115 x dHOPEMTasd |
| **FL4003** | *Zdk1-SctQ, mCherry-SctL* | dHOPEMTasd | pAD612 x ADTM4521 |
| **FL4004** | *SspB_Nano-mCherry-SctQ* | dHOPEMTasd | pFL117 x dHOPEMTasd |
| **FL4005** | *SspB_Nano-SctQ, mCherry-SctL* | dHOPEMTasd | pFL118 x ADTM4521 |

#### Suppl. Table 2: *Yersinia enterocolitica* strains used in this study

* x = homologous recombination between mutator-plasmid and host-strain, recombination leads to an allelic exchange of the native gene and the mutated gene[12].

| **Name** (Reference) | **Genotype** | **Primers for amplification** | **Template for Insert** | **System** |
|---|---|---|---|---|
| **pFL100** (this work) | *pBAD::TMH-FLAG-(L1)-mCherry-(L2)-LOV2* | AD638/704/705 | p81041* | LOV |
| **pFL101** (this work) | *pACYC184::Zdk1-(L4)-EGFP* | AD698/699/ 700/701 | P81010*/ pAD301 | LOV |
| **pFL104** (this work) | *pACYC184::Zdk1-(L4)-mCherry* | AD638/699/ 721/722 | p81010*/ pAD304 | LOV |
| **pFL107** (this work) | *pBAD::TMH-FLAG-(L1)-mCherry-(L3)-iLID* | AD638/706/707/ 732/733 | p60408*/ pAD304 | iLID |
| **pFL108** (this work) | *pBAD::TMH-FLAG-(L1)-iLID* | AD638/733/734 | p60408* | iLID |
| **pFL109** (this work) | *pACYC184::SspB_Nano-(L4)-mCherry* | AD721/722/ 735/736 | p60409*/ pAD304 | iLID |
| **pFL111** (this work) | *pBAD::Zdk1-(L4)-mCherry* | AD759/768 | pFL104 | LOV |
| **pFL113** (this work) | *pBAD::SspB_Nano-(L4)-mCherry* | AD762/768 | pFL109 | iLID |
| **pFL114** (this work) | *pBAD::SspB_Nano* | AD762/769 | pFL109 | iLID |
| **pFL115** (this work) | *pKNG101::Zdk1-(L4)-mCherry-SctQ* | ** | pFL111 | LOV |
| **pFL117** (this work) | *pKNG101::SspB_Nano-(L4)-mCherry-SctQ* | ** | pFL113 | iLID |
| **pFL118** (this work) | *pKNG101::SspB_Nano-SctQ* | ** | pFL114 | iLID |
| **pFL126** (this work) | *pACYC184::TMH-FLAG-(L1)-LOV2*$_{V416L}$ | AD921/903 | pAD610 | LOV |
| **pFL127** (this work) | *pACYC184::TMH-FLAG-(L1)-iLID* | AD921/904 | pFL108 | iLID |
| **pFL133** (this work) | *pBAD::YopE*$_{1\text{-}53}$*-Nanoluc-FLAG* | | pAD681 | |
| **pAD304** (ref. [3]) | *pUC19-mCherry* | | | |
| **pAD608** (this work) | *pBAD::TMH-FLAG-(L1)-LOV2* | AD638/639/640 | p81041* | LOV |
| **pAD610** (this work) | *pBAD::TMH-FLAG-(L1)-LOV2*$_{V416L}$ | | | LOV |
| **pAD612** (this work) | *pKNG101::Zdk1-SctQ* | ** | pAD611 | LOV |
| **pBMD028** (this work) | *pBAD::YopE*$_{1\text{-}53}$ | AD894/895 | pYV(MRS40) | |
| **pBMD040** (this work) | *pBAD::YopE*$_{1\text{-}53}$*-*$\beta$*-lactamase* | ** | pNL1.1 (Promega) /pBMD028 | |

#### Suppl. Table 3: Plasmids used in this study

Plasmids with corresponding properties that were designed and/or used in this work. Anchor proteins were targeted to the bacterial IM by addition of an optimized TMH based on the N-terminal TMH of the *Escherichia coli* TatA protein[13], an integral component of the Tat export system[14]. A high expression ratio of the anchor to bait protein was reported to be a prerequisite for complete binding of the bait to the anchor[15]. We therefore expressed the membrane anchor constructs from the inducible medium-high copy expression vector pBAD-His/B, and the cytosolic bait fusions from a compatible low copy constitutive expression vector, pACYC184. * Addgene code; ** Restriction and ligation only; L1 – GAGG linker; L2 – GSGS linker; L3 – GAGGGAGG linker; L4 – GGSGGSGG linker.

| Primer | Sequence from 5' to 3' | Reference |
|---|---|---|
| AD638 | GGTCTCC**ATG**GGTGGTATCAGTATTTGGCAGTTATTGATTATTGCCGTCATCGTTGTACTGCTTGTCCTATTTGGCACCAAAAAGCTCGGCTCCGACTACAAGGACGACGATGATAAGGGTGGAGCAGGT | this work |
| AD639 | GACTACAAGGACGACGATGATAAGGGTGGAGCAGGTGGATCCTTGGCTACTACACTTGA | this work |
| AD640 | GACTGAATTCGCAAGCTT*TTA*AAGTTCTTTTG | this work |
| AD698 | GACTGGATCCT*TGACTGA*ATGGTGGATAACAAATTCAATAAAGAAAAGA | this work |
| AD699 | ACCACCAGAGCCGCCCGACCCACCAGAACCACCTTTTGGGGCCT | this work |
| AD700 | GGTGGGTCGGGCGGCTCTGGTGGTGGTGCTGGCGTGAGCAAG | this work |
| AD701 | GATCGTCGACTTACTTGTACAGCTCGTCCATGC | this work |
| AD704 | GACGATGATAAGGGTGGAGCAGGTGTGAGCAAGGGCGAGGAG | this work |
| AD705 | GACTGAATTCGCAAGCTTTTAAAGTTCTTTTG | this work |
| AD706 | GACGATGATAAGGGTGGAGCAGGTGTGAGCAAGGGCGAGGAG | this work |
| AD707 | TCCACCTGCTCCACCACCAGCGCCCTTGTACAG | this work |
| AD721 | GGTGGGTCGGGCGGCTCTGGTGGTGTGAGCAAGGGCGAGGAG | this work |
| AD722 | GATCGTCGACTTACTTGTACAGCTCGTCCATGC | this work |
| AD732 | GGCGCTGGTGGTGGAGCAGGTGGAGGATCCGGGGAGTTTCTGG | this work |
| AD733 | GACTGAATTCTCAGCTAATTAAGCTTTTAAAAGT | this work |
| AD734 | GACGATGATAAGGGTGGAGCAGGTGGATCCGGGGAGTTTCTGG | this work |
| AD735 | GACTGGATCCT*TGACTGA*ATGAGCTCCCCGAAACGCCC | this work |
| AD736 | ACCACCAGAGCCGCCCGACCCACCACCAATATTCAGCTCGTCAT | this work |
| AD759 | GACTAGATCTGGCGCAGGTGTGGATAACAAATTCAATAAAGAAAAGA | this work |
| AD762 | GACTAGATCTGGCGCAGGTAGCTCCCCGAAACGCCCTAA | this work |
| AD768 | GACTGAATTCACCTGCGCCCTTGTACAGCTCGTCCATGC | this work |
| AD769 | GACTGAATTCACCTGCGCCACCAATATTCAGCTCGTCATAGA | this work |
| AD894 | GATCTC**ATGA**AAATATCATCATTTATTTCTACATCACTGC | this work |
| AD895 | GATCGAATTCGCGCAGATCTTCCGCCGGAACCCTGAGGGCTTTCAGTGC | this work |
| AD903 | GACTGTCGACGCAAGCTT*TTA*AAGTTCTTTTG | this work |
| AD904 | GACTGTCGACTCAGCTAATTAAGCTTTTAAAAGT | this work |
| AD921 | GACTAGATCT*TTGACTGA*ATGGGTGGTATCAGTATTTGGC | this work |
| AD933 | GACTTC***ATG****AACTTATCATTAAGCGATCTTCATCGTCAGGTATCTCGATTGGTGCAG*GGAGGTAGATCTGGCGCAGGTGTCTTCACACTCGAAGATTTCGTT | this work |
| AD934 | GACTAAGCTTTTAGAATTCGCCTGCACC*CTTATCATCGTCGTCCTTGTAGTC*ACCTCCCGCCAGAATGCGTTCGCA | this work |

**Suppl. Table 4: Primers used in this study**

Red, restriction sites; blue, linker; purple/bold = start/stop sites.

| **Fluorescence ratio** | **Average** | **St.dev.** |
|---|---|---|
| TMH-FLAG-mCherry-LOV2 / Zdk1-mCherry-SctQ | 1,955293 | 0,244407 |
| TMH-FLAG-LOV2$_{V416L}$ / TMH-FLAG-LOV2 | 1,232724 | 0,926356 |
| **anchor / bait** = TMH-FLAG-LOV2$_{V416L}$ / Zdk1-SctQ | 2,410336 | 1,836185 |

**Suppl. Table 5: Calculation of ancor/bait ratios**

Experimentally determined (blue) and calculated (orange) expression ratios of anchor and bait proteins for the LITESEC-act2 system; n=3.
